# White matter aberrations and age-related trajectories in patients with schizophrenia and bipolar disorder revealed by diffusion tensor imaging

**DOI:** 10.1101/395889

**Authors:** Siren Tønnesen, Tobias Kaufmann, Nhat Trung Doan, Dag Alnæs, Aldo Córdova-Palomera, Dennis van der Meer, Jaroslav Rokicki, Torgeir Moberget, Tiril P. Gurholt, Unn K. Haukvik, Torill Ueland, Trine Vik Lagerberg, Ingrid Agartz, Ole A. Andreassen, Lars T. Westlye

## Abstract

Supported by histological and genetic evidence implicating myelin, neuroinflammation and oligodendrocyte dysfunction in schizophrenia spectrum disorders (SZ), diffusion tensor imaging (DTI) studies have consistently shown white matter (WM) abnormalities when compared to healthy controls (HC). The diagnostic specificity remains unclear, with bipolar disorders (BD) frequently conceptualized as a less severe clinical manifestation along a psychotic spectrum. Further, the age-related dynamics and possible sex differences of WM abnormalities in SZ and BD are currently understudied.

Using tract-based spatial statistics (TBSS) we compared DTI-based microstructural indices between SZ (n=128), BD (n=61), and HC (n=293). We tested for age-by-group and sex-by-group interactions, computed effect sizes within different age-bins and within genders.

TBSS revealed global reductions in fractional anisotropy (FA) and increases in radial (RD) diffusivity in SZ compared to HC, with strongest effects in the body and splenium of the corpus callosum, and lower FA in SZ compared to BD in right inferior longitudinal fasciculus and right inferior fronto-occipital fasciculus, and no significant differences between BD and HC. The results were not strongly dependent on age or sex. Despite lack of significant group-by-age interactions, a sliding-window approach supported widespread WM involvement in SZ with most profound differences in FA from the late 20s.

## Introduction

Schizophrenia (SZ) is a major cause of years lived with disability^1^, yet the pathological mechanisms underlying the diverse manifestations of this disorder remain unclear^2^. In line with the notion that the symptoms partly arise from abnormal brain connectivity and functional integration of brain processes^3-5^, histopathological and genetic investigations have implicated lipid homeostasis, neuroinflammation, myelin and oligodendrocyte abnormalities^6-11^. Furthermore, brain imaging has consistently indicated anatomically widespread white matter (WM) microstructural abnormalities using diffusion tensor imaging (DTI)^12-15^. A recent meta-analysis across 29 cohorts of the ENIGMA consortium, including the current, documented significantly lower fractional anisotropy (FA) in patients with SZ (n=1963) compared to healthy controls (n=2359) in 20 of 25 regions of interest, and no significant associations with age of onset and medication status^16^.

WM abnormalities have been reported across a wide range of clinical traits and disorders^17-22^, and the diagnostic specificity remains unknown. Current diagnostic nosology treats SZ and bipolar spectrum disorder (BD) as independent categories, but genetic^23^, clinical and neuropsychological^24,25^ evidence suggest partly overlapping pathophysiology and clinical manifestation, with higher symptom burden, poorer function and worse outcome in SZ^26^. Including both patients with SZ and BD in the same analysis is vital for probing common and distinct etiological mechanisms across the psychosis spectrum. While DTI aberrations have consistently been documented in BD^27-31^, the existing studies that have included both groups have not provided conclusive evidence of marked group differences between BD and SZ^32-39^.

SZ have been conceptualized as a neurodevelopmental disorder^40,41^ and deficient myelination during adolescence has been included among the core features of the prodromal phase^42^ with WM aberrations present before disease onset^43,44^. Along with evidence of accelerated brain changes in adult SZ^45-47^, neurodevelopmental theories strongly support the need for a dynamic lifespan perspective. The magnitude and modulators of group differences vary across life, e.g. manifested as delayed developmental trajectories^48^ and progressive aging-related changes in adulthood^49^. Moreover, whereas the largest meta-analysis to date revealed no significant group by sex interactions, effect sizes for female patients were significantly larger compared to the effect sizes for males for global FA^16^ (but see another recent review and meta-analysis which found no significant sex-related differences in effect sizes when comparing patients and controls within males and females, respectively^50^). Although the evidence of strong modulating effects of sex on WM abnormalities in severe mental disorders is lacking, sexual dimorphisms in brain biology and clinical expression warrant further studies on possible sex by diagnosis interactions on the human brain.

In order to address these unresolved issues, our main aim was to compare several DTI indices across the brain between patients diagnosed with a SZ spectrum disorder (n=128), BD (n=61), and HC (n=293), both within and across males and females, using tract-based spatial statistics (TBSS)^51^. Based on the literature we expected widespread WM microstructural alterations in SZ compared to HC, in particular lower FA. Additionally, based on clinical severity, we anticipated moderate group differences between BD and HC, with BD showing less distinct and distributed abnormalities. To comply with a dynamic age-variant perspective, we tested for group by age interactions and compared effect sizes within age cohorts using a sliding window technique^52^, and also tested for group by sex interactions and compared effect sizes within females and males, respectively. DTI based indices of WM microstructure are highly sensitive to differences in data quality, e.g. due to subject motion, which may bias the results^53-55^. Since previous studies have often failed to report quality control (QC) measures or simply omitted systematic QC altogether, it is unknown if reported group effects are biased due to differences in data quality. Hence, we employed a stringent multi-step exclusion protocol based on quantitative quality assessment, and compared groups at different levels to assess the relevance.

## Results

### Demographics and clinical characteristics

Table 1 summarizes demographics and clinical characteristics. There was a significant main effect of group on age (F=4.2, p=.016), education (F=34.1, p<.001) and IQ (F=37.3, p<.001), with higher age in HC compared to SZ (Supplementary Fig. S1), longer education and higher IQ (HC>BD>SZ). Compared to BD, SZ had higher symptom severity as measured by The Positive and Negative Syndrome Scale (PANSS) total (t=8.2, p<.001), positive (t=7.6, p<.001), negative (t=6.4, p<.001), and disorganized (t=4.2, p<.001) sub-scales, and the split version of Global Assessment of Functioning Scale split version (GAF)^56^ with GAF function (t=-5.5, p<.001) and GAF symptom (t=-7.4, p<.001).

**Table 1.**
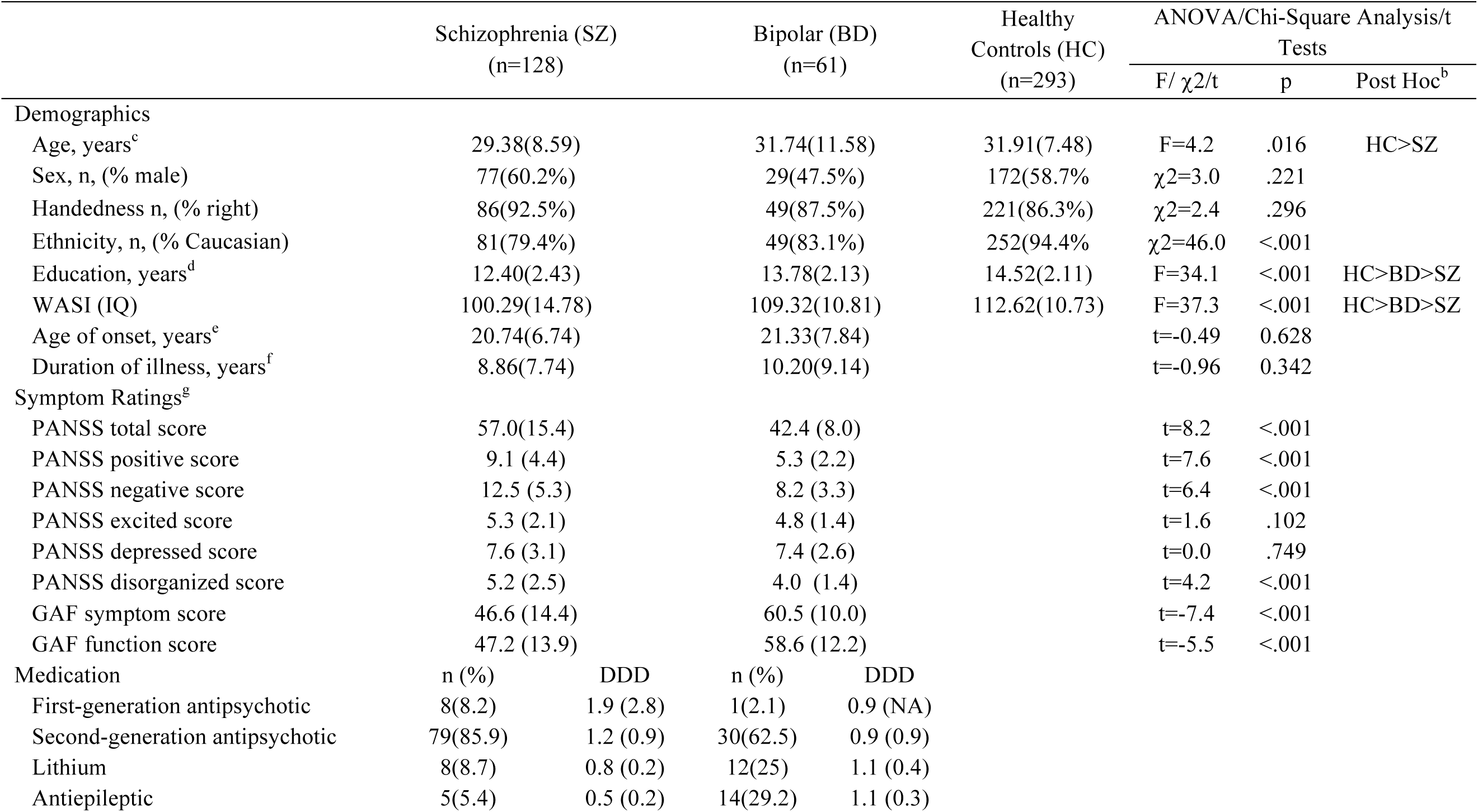
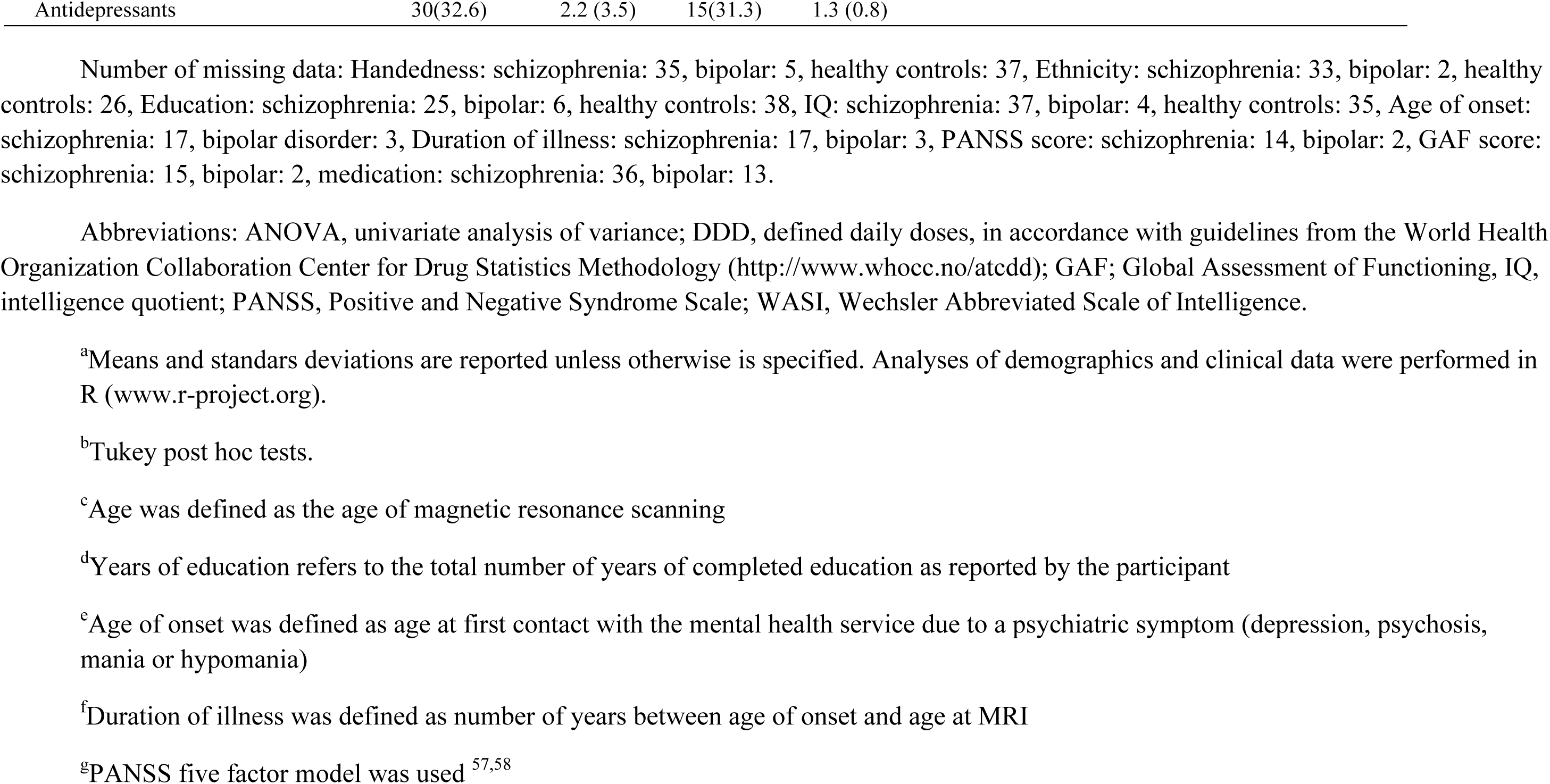
Demographic and clinical Data^a^

|  | Schizophrenia (SZ)<br>(n=128) | Bipolar (BD)<br>(n=61) | Healthy<br>Controls (HC)<br>(n=293) | ANOVA/Chi-Square Analysis/t<br>Tests |  |  |
| --- | --- | --- | --- | --- | --- | --- |
| | | | | F/ $\chi^2$/t | p | Post Hoc<sup>b</sup> |
| Demographics |  |  |  |  |  |  |
| Age, years<sup>c</sup> | 29.38(8.59) | 31.74(11.58) | 31.91(7.48) | F=4.2 | .016 | HC>SZ |
| Sex, n, (% male) | 77(60.2%) | 29(47.5%) | 172(58.7%) | $\chi^2$=3.0 | .221 | |
| Handedness n, (% right) | 86(92.5%) | 49(87.5%) | 221(86.3%) | $\chi^2$=2.4 | .296 | |
| Ethnicity, n, (% Caucasian) | 81(79.4%) | 49(83.1%) | 252(94.4%) | $\chi^2$=46.0 | <.001 | |
| Education, years<sup>d</sup> | 12.40(2.43) | 13.78(2.13) | 14.52(2.11) | F=34.1 | <.001 | HC>BD>SZ |
| WASI (IQ) | 100.29(14.78) | 109.32(10.81) | 112.62(10.73) | F=37.3 | <.001 | HC>BD>SZ |
| Age of onset, years<sup>c</sup> | 20.74(6.74) | 21.33(7.84) |  | t=-0.49 | 0.628 |  |
| Duration of illness, years<sup>f</sup> | 8.86(7.74) | 10.20(9.14) |  | t=-0.96 | 0.342 |  |
| Symptom Ratings<sup>g</sup> |  |  |  |  |  |  |
| PANSS total score | 57.0(15.4) | 42.4 (8.0) |  | t=8.2 | <.001 |  |
| PANSS positive score | 9.1 (4.4) | 5.3 (2.2) |  | t=7.6 | <.001 |  |
| PANSS negative score | 12.5 (5.3) | 8.2 (3.3) |  | t=6.4 | <.001 |  |
| PANSS excited score | 5.3 (2.1) | 4.8 (1.4) |  | t=1.6 | .102 |  |
| PANSS depressed score | 7.6 (3.1) | 7.4 (2.6) |  | t=0.0 | .749 |  |
| PANSS disorganized score | 5.2 (2.5) | 4.0 (1.4) |  | t=4.2 | <.001 |  |
| GAF symptom score | 46.6 (14.4) | 60.5 (10.0) |  | t=-7.4 | <.001 |  |
| GAF function score | 47.2 (13.9) | 58.6 (12.2) |  | t=-5.5 | <.001 |  |
| Medication | n (%) | DDD | n (%) | DDD |  |  |
| First-generation antipsychotic | 8(8.2) | 1.9 (2.8) | 1(2.1) | 0.9 (NA) |  |  |
| Second-generation antipsychotic | 79(85.9) | 1.2 (0.9) | 30(62.5) | 0.9 (0.9) |  |  |
| Lithium | 8(8.7) | 0.8 (0.2) | 12(25) | 1.1 (0.4) |  |  |
| Antiepileptic | 5(5.4) | 0.5 (0.2) | 14(29.2) | 1.1 (0.3) |  |  |

|  |  |  |  |  |
| --- | --- | --- | --- | --- |
| Antidepressants | 30(32.6) | 2.2 (3.5) | 15(31.3) | 1.3 (0.8) |

Number of missing data: Handedness: schizophrenia: 35, bipolar: 5, healthy controls: 37, Ethnicity: schizophrenia: 33, bipolar: 2, healthy controls: 26, Education: schizophrenia: 25, bipolar: 6, healthy controls: 38, IQ: schizophrenia: 37, bipolar: 4, healthy controls: 35, Age of onset: schizophrenia: 17, bipolar disorder: 3, Duration of illness: schizophrenia: 17, bipolar: 3, PANSS score: schizophrenia: 14, bipolar: 2, GAF score: schizophrenia: 15, bipolar: 2, medication: schizophrenia: 36, bipolar: 13.
Abbreviations: ANOVA, univariate analysis of variance; DDD, defined daily doses, in accordance with guidelines from the World Health Organization Collaboration Center for Drug Statistics Methodology (http://www.whocc.no/atcdd); GAF; Global Assessment of Functioning, IQ, intelligence quotient; PANSS, Positive and Negative Syndrome Scale; WASI, Wechsler Abbreviated Scale of Intelligence.
<sup>a</sup>Means and standards deviations are reported unless otherwise is specified. Analyses of demographics and clinical data were performed in R (www.r-project.org).
<sup>b</sup>Tukey post hoc tests.
<sup>c</sup>Age was defined as the age of magnetic resonance scanning
<sup>d</sup>Years of education refers to the total number of years of completed education as reported by the participant
<sup>e</sup>Age of onset was defined as age at first contact with the mental health service due to a psychiatric symptom (depression, psychosis, mania or hypomania)
<sup>f</sup>Duration of illness was defined as number of years between age of onset and age at MRI
<sup>g</sup>PANSS five factor model was used <sup>57,58</sup>

### Main effects of diagnostic groups on DTI

Figure 1 and summarize results from voxelwise analyses testing for main effects of group on the DTI indices. We found significant and widespread main effects of group on FA and radial diffusivity (RD), including the corpus callosum, superior longitudinal fasciculus, fornix, cingulum, forceps major and inferior fronto-occipital fasciculus. Pairwise comparisons revealed widespread FA reductions and RD increases in SZ compared to HC, and FA reductions in SZ compared to BD. No other group comparisons yielded significant effects.

**Figure 1.**
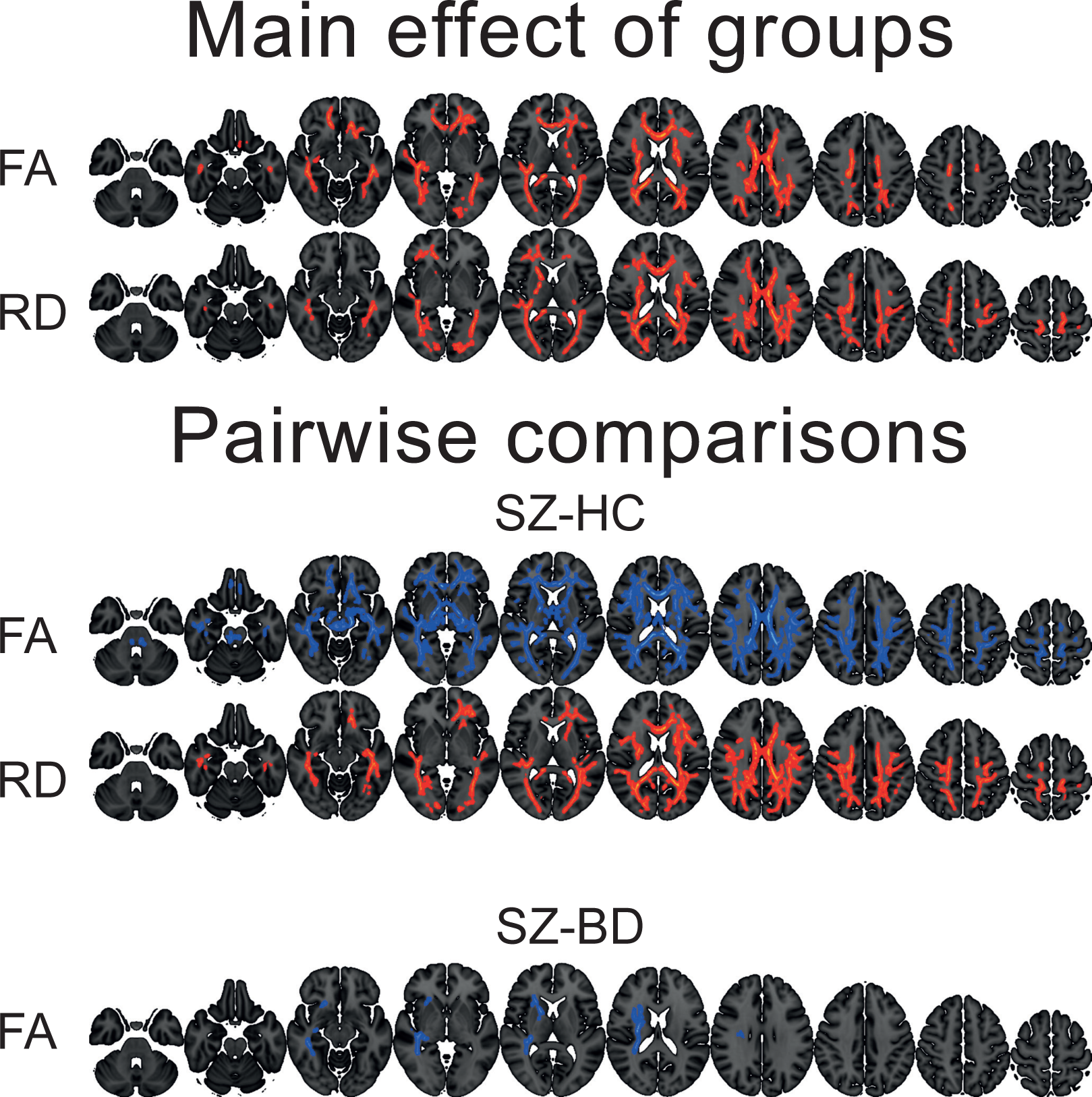
Colored voxels show significantly decreased (blue) and increased (red) DTI-indices in SZ patients relative to HC and BD. Group differences are thresholded at *p* <.05 (two-tailed) after permutation testing using threshold free cluster enhancement (TFCE). Note that the white matter skeleton has been slightly thickened to aid visualisation.

### Global DTI measures and sliding window approach

Figure 2 shows mean skeleton DTI values plotted as a function of age and group. Table 2 summarizes the results from linear models accounting for age, age^2,^ sex and diagnosis. We found significant main effects of group, sex and age on FA. There was no significant group by age or group by age^2 interaction; therefore the model was run without the interaction terms. Pairwise comparisons revealed lower FA in SZ compared to HC, and lower FA in SZ compared to BD. Females showed significantly lower FA compared to males. Figure 2, Supplemental Fig. S2 and Table 3 summarize results from the bootstrapped age fitting procedure, yielding estimates of the mean and standard deviation of age at maximum FA and minimum RD, MD and axial diffusivity (AD) within groups. The overlap in confidence intervals (Table 3) and the comparison against empirical null distributions generated using permutation testing (see Methods and Supplemental Fig. S2) revealed no significant between-group differences in age at maximum (FA) or minimum (RD, MD, and AD). Supplemental Fig. S3-S6 summarize the bootstrapped age fitting procedure for regions-of-interest (ROI). Briefly, the majority of ROIs show a consistent pattern with early peak in FA in SZ compared to BD and HC. However, the results from left and right cingulum have a more intricate pattern with BD reporting an older age peak. For RD, AD and MD the trend is less consistent and more complex.

**Table 3.**
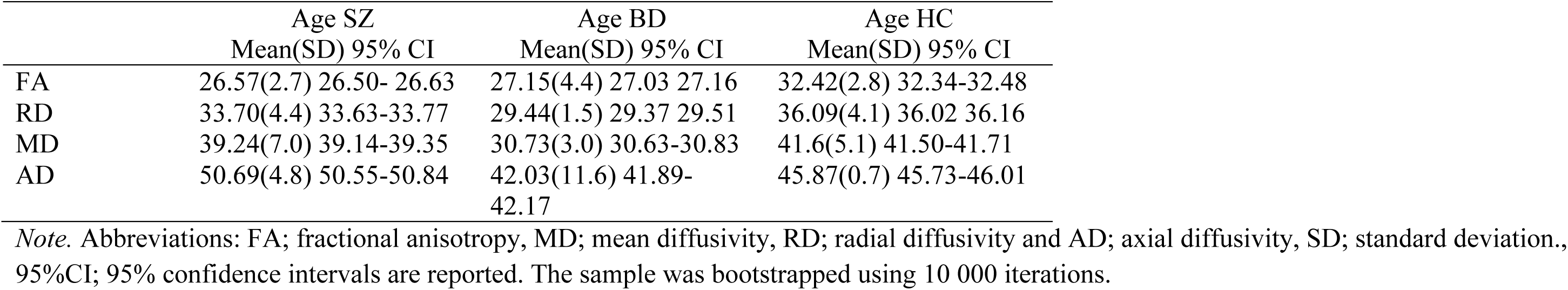
Age where maximum FA or minimum RD, MD or AD were reached

**Table 2.** Mean skeleton DTI metrics within groups

| | SZ<br>Mean(SD)<br>95%CI | BD<br>Mean(SD)<br>95%CI | HC<br>Mean(SD)<br>95%CI | F | $p^{(a)}$ | Pairwise<br>comparison | Cohens d<br>HC/SZ,HC/BD,<br>BD/SZ | Age<sup>2</sup><br>t | Age<sup>2</sup><br>p | Age<br>t | Age<br>p | Sex<br>t | Sex<br>p |
| --- | --- | --- | --- | --- | --- | --- | --- | --- | --- | --- | --- | --- | --- |
| FA | 0.452(0.01)<br>0.450-0.455 | 0.455(.01)<br>0.451-0.458 | 0.458(0.01)<br>0.457-0.460 | 7.02 | <0.001* | HC>SZ,<br>BD>SZ | 0.35, 0.03, 0.21 | -3.97 | <0.001 | 3.49 | <0.001 | 4.98 | <0.001 |
| MD | 782.04(22.01)<br>778.19-785.89 | 782.11(20.16)<br>776.95-787.27 | 776.71(19.22)<br>774.50-778.92 | 0.82 | 0.437 |  | -0.11, -0.06, -<br>0.03 | 5.06 | <0.001 | -5.63 | <0.001 | -1.16 | 0.247 |
| RD | 571.82(22.04)<br>567.97-575.68 | 571.06(21.65)<br>565.52-576.61 | 564.49(20.34)<br>562.15-566.83 | 2.96 | 0.052 |  | -0.23, -0.07, -<br>0.09 | 5.21 | <0.001 | -5.29 | <0.001 | -2.24 | 0.025 |
| AD | 1202.49(27.94)<br>1197.60-1207.38 | 1204.20(22.96)<br>1198.32-1210.08 | 1201.16(22.08)<br>1198.62-1203.69 | 0.71 | 0.492 |  | 0.11, -0.01, 0.09 | 3.69 | <0.001 | -5.08 | <0.001 | 1.11 | 0.273 |

*Note.* Abbreviations: FA; fractional anisotropy, MD; mean diffusivity, RD; radial diffusivity and AD; axial diffusivity. MD,RD,AD multiplied by 1 000 000 to preserve precision. The p-values are not corrected for multiple testing. There was no significant group by age interaction or group by age<sup>2</sup> interaction; therefore the model was run without these interaction terms. The effect size represent the group by age interaction in the age squared model ( $x \sim \text{group} + \text{age} + \text{age}^2 + \text{sex} + \text{group}:\text{age}$ ). \* Bonferroni corrected $p < .05$

**Figure 2.**
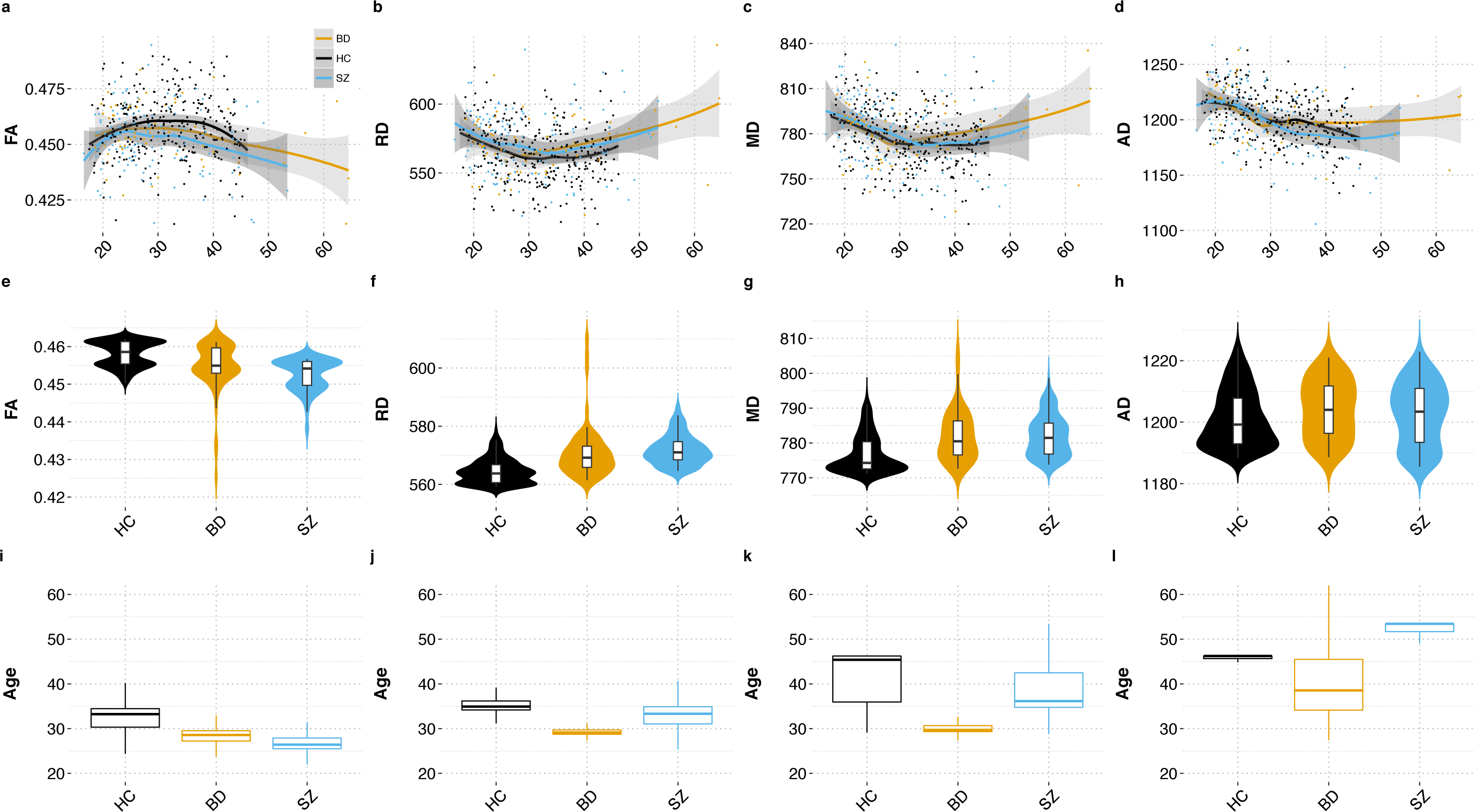
Plots a-d: Mean skeleton DTI values plotted as a function of age and group (HC = healthy controls, BD = bipolar disorders, SZ = schizophrenia spectrum disorders). Plots e-h: Violin plot depicting the fitted values for each group. Plots i-l: Uncertainty estimates of the age within each group when maximum FA, minimum RD, MD and AD are reached from a bootstrap procedure with 10 000 resamples.

Effect sizes within different age-bins from the sliding-window technique are presented in Supplementary Fig. S7. For FA, MD, RD, effect sizes for HC vs. SZ increased until the late 20s. Effect sizes for SZ vs. BD showed a similar pattern for FA, and more complex non-linear associations for MD, RD and AD. Effect sizes for HC vs. BD for FA straddled around 0 throughout the sampled age range. For RD, MD, and AD the effect sizes showed more complex non-linear associations.

### Sex related differences

Mean skeleton and ROI analyses revealed no significant sex-by-diagnosis interaction effects on DTI WM metrics. Supplementary Table S1 shows results from group comparisons within females and males, respectively. Briefly, the analysis revealed main effects of group on FA both in males (F=3.36, p=.036) and females (F=3.99, p=.02). Pairwise comparisons revealed lower FA in SZ compared to HC in both sexes and lower FA in SZ compared to BD in females.

### Associations with symptom domains

Mean skeleton and ROI analyses revealed no significant associations with GAF and PANSS domain scores across patient groups. Global and ROI-based t- and p-statistics are summarized in Supplementary Table S2 and Supplementary Fig. S8, respectively.

### Effects of quality control

Visual inspection of the datasets with QC summary z-score below -2.5 (n=35, see below) indicated no clear reason for exclusion. Therefore, the main analyses were run on the entire dataset, but we also present results using varying QC levels. Figure 3 summarizes the effects of QC on the mean skeleton data. Effect sizes for HC vs. SZ and BD vs. SZ increased with QC stringency for all metrics except AD, which showed a more complex pattern. The effect size for HC vs. BD remained relative unchanged as a function of QC. Voxelwise analysis revealed highly similar patterns as those obtained using the full sample (Fig. 1 and Supplementary Fig. S9).

**Figure 3.**
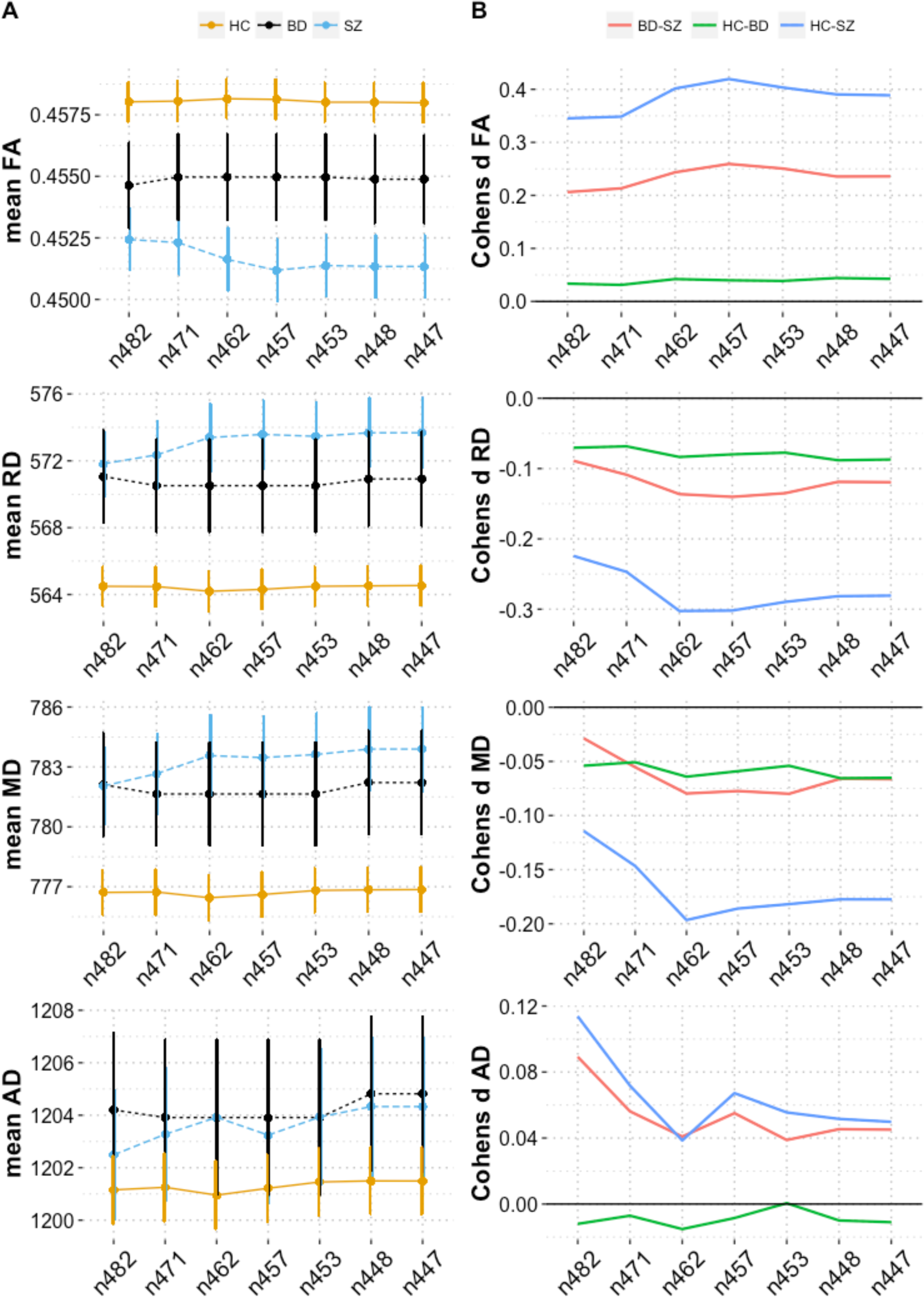
Mean of skeleton DTI metrics plotted across quality control subgroup analyses (A) and Cohens d for pairwise comparisons across quality control subgroup analyses (B). The labels on the x-axes reflect the number of participants in each analysis. The error bars of part A represent the standard error of the mean.

### Subgroup and age restricted analyses

Density plots showing the distribution of the four DTI metrics for each of the subgroups within each diagnostic group are presented in Supplementary Fig. S10. The results from the subgroup analyses are presented in Supplementary Table S3, while the age restricted analyses are presented in Supplementary Table S4. Briefly, but not limited to, for the diagnostic subgroups there were significant differences between psychosis not otherwise specified (PNOS) and a strict SZ diagnosis for global MD (*p*=0.011, uncorrected) and global AD (*p*=0.002, uncorrected). A similar pattern was observed for the BD subgroups, with significant difference between BDI and BDII for MD (*p*=0.035, uncorrected) and AD (*p*=0.035, uncorrected). For the age-restricted analyses (55 years and younger) we observed main effect of group for FA only, with pairwise comparisons indicating lower global FA in SZ compared to HC.

### ROI analyses

Figure 4 and Table 4 summarize the ROI results. Most ROIs showed main effects of group for FA (η ps^2^: 0.001-0.029), with strongest effects in the body (BCC) and splenium (SCC) of the corpus callosum and forceps major. We found substantial effects of group in 7 and 1 of the 23 ROIs in RD and AD. We found a nominal significant (p<.05, uncorrected) age by group interactions for MD in the BCC (η ps^2^<0.013, *p*=0.044), indicating larger group differences with increasing age. Since no age by group interactions remained after corrections for multiple comparisons, all main effects and results from pairwise comparisons were computed without the interaction term in the models.

**Table 4.**
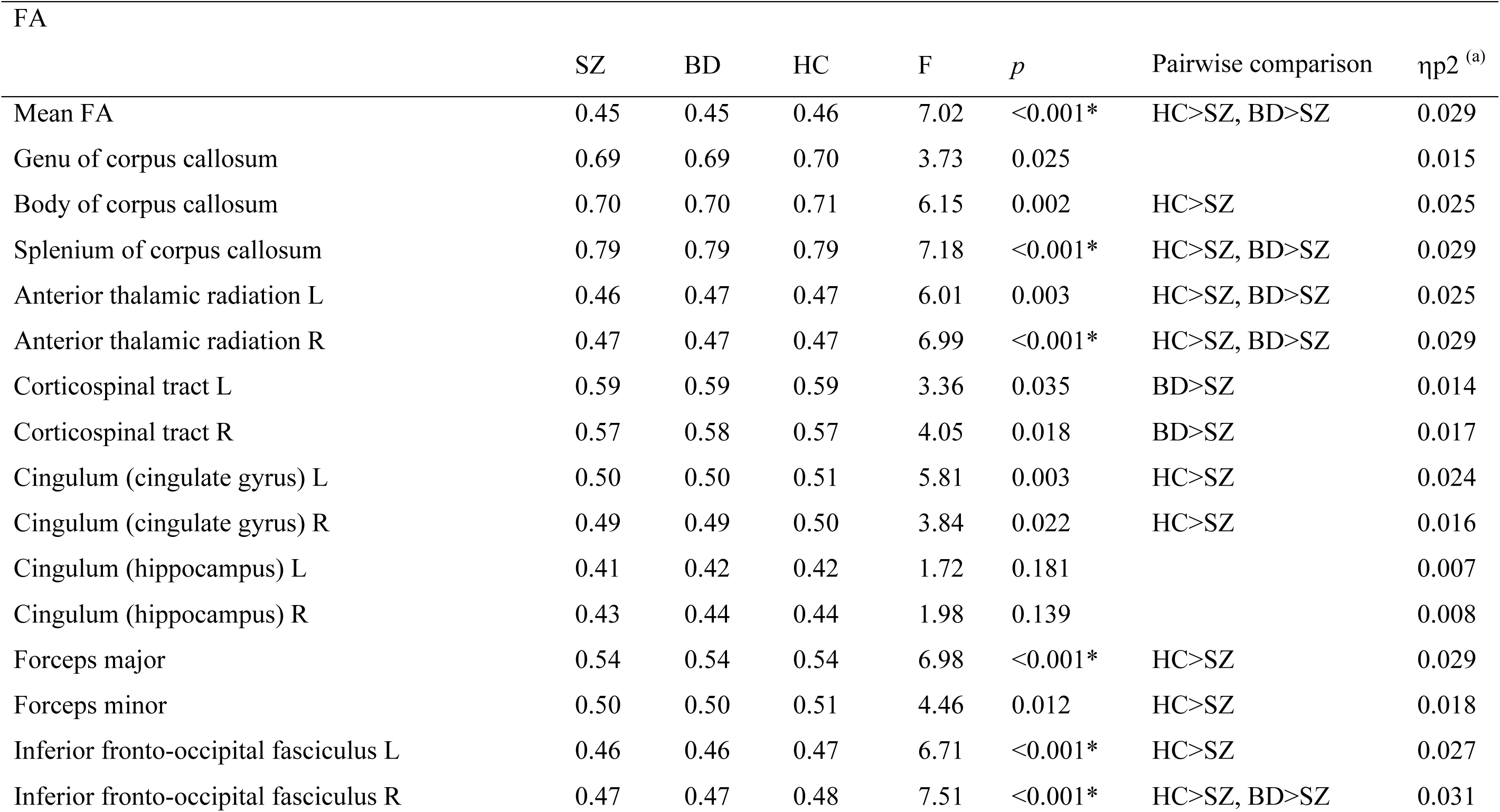
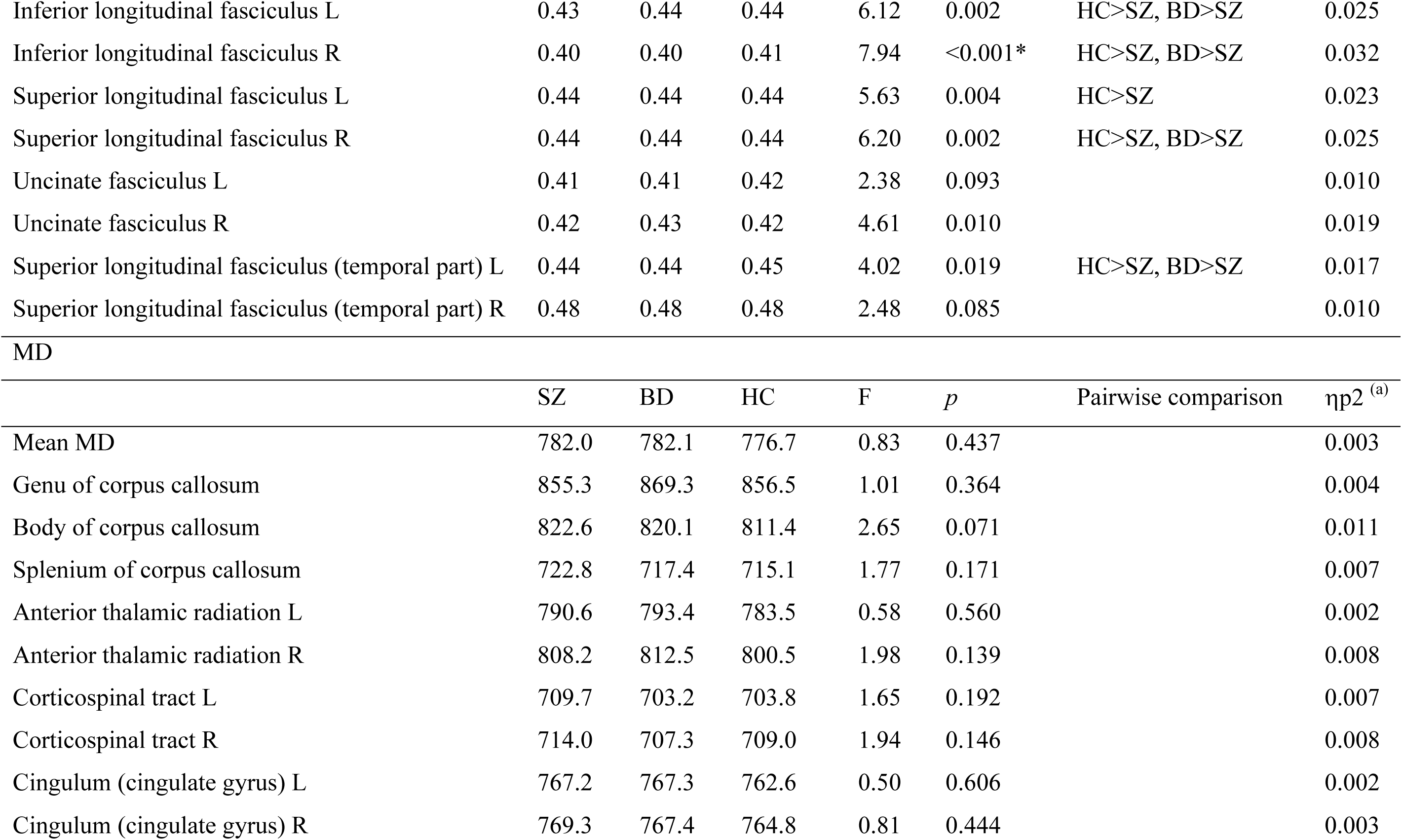
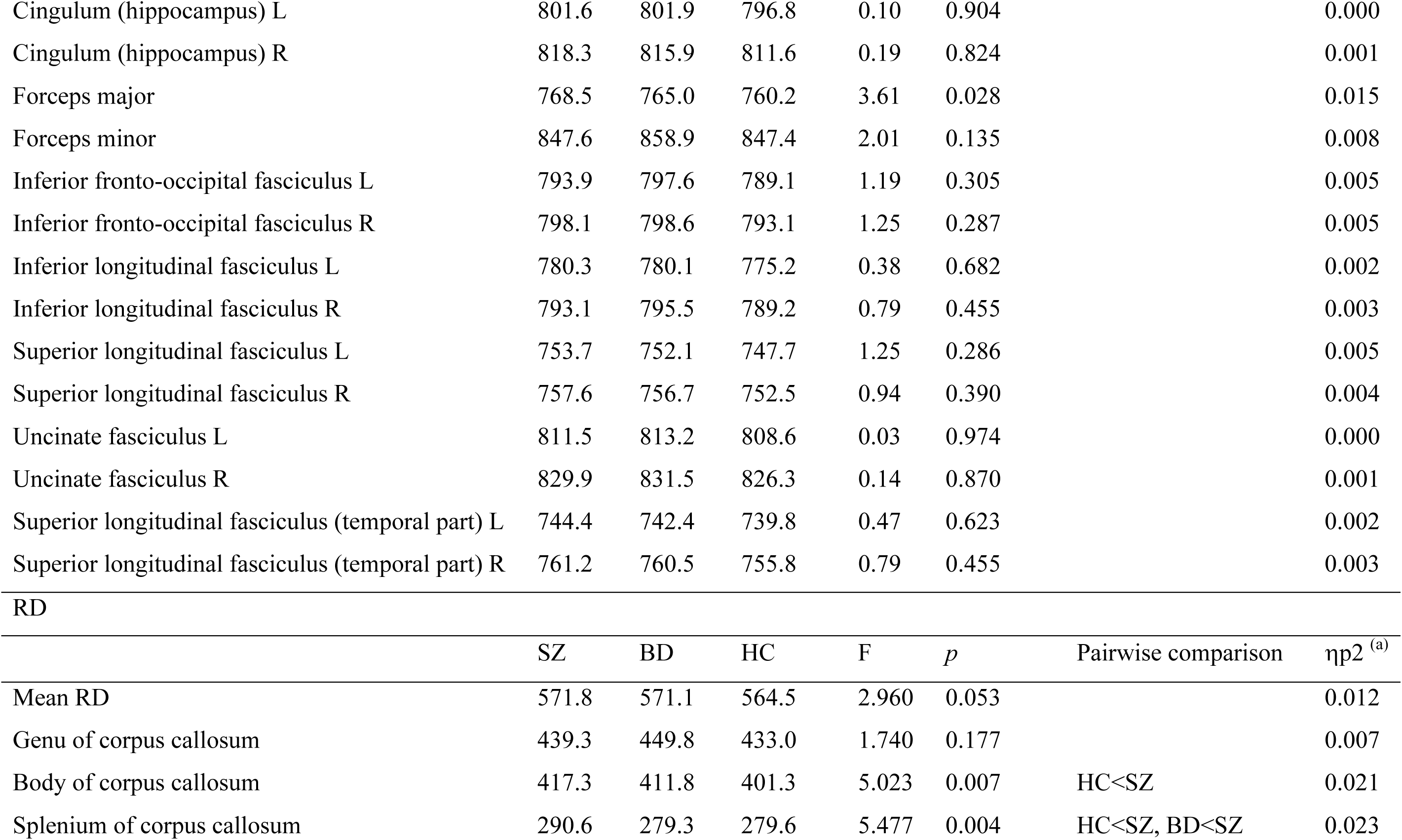
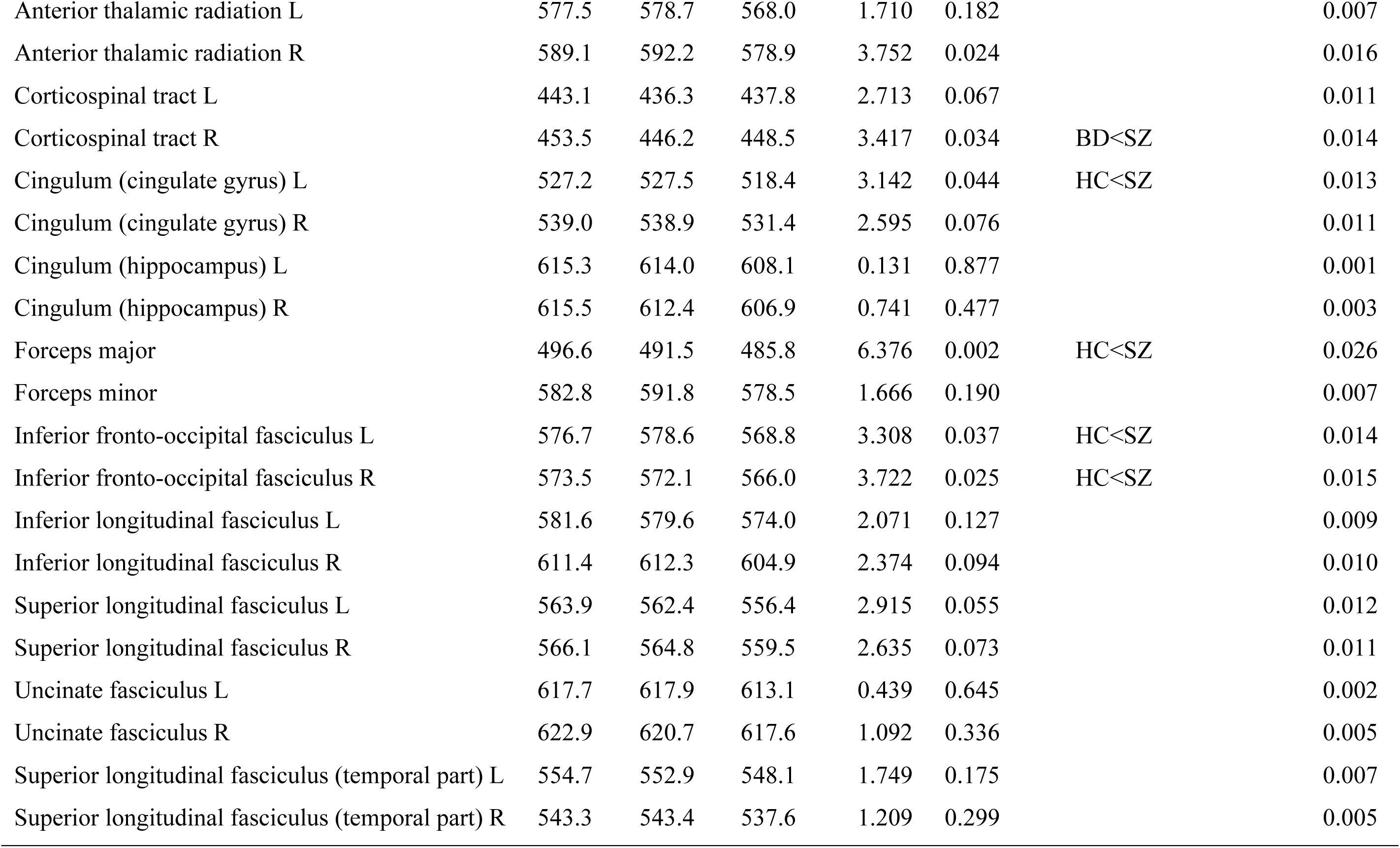
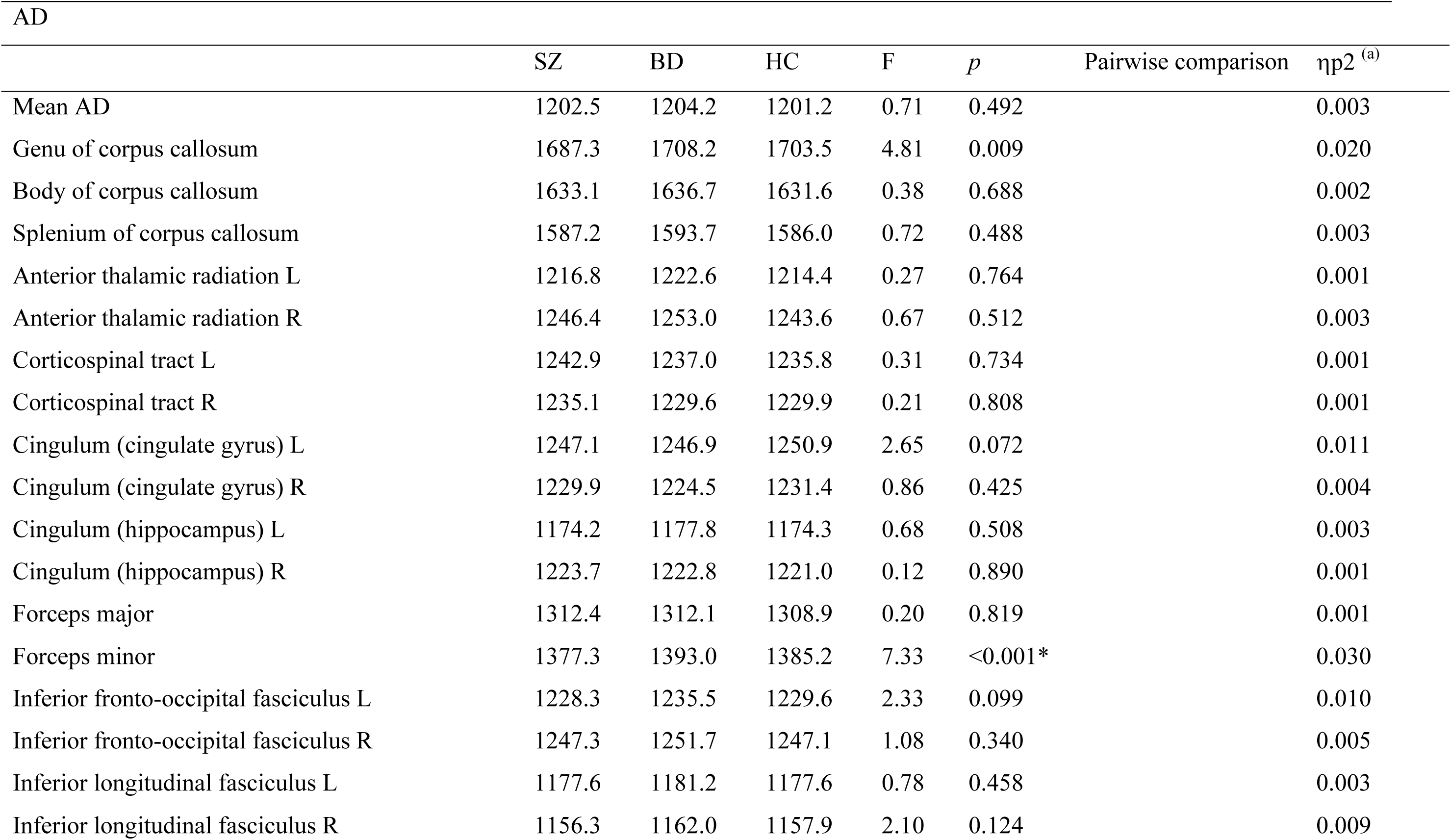
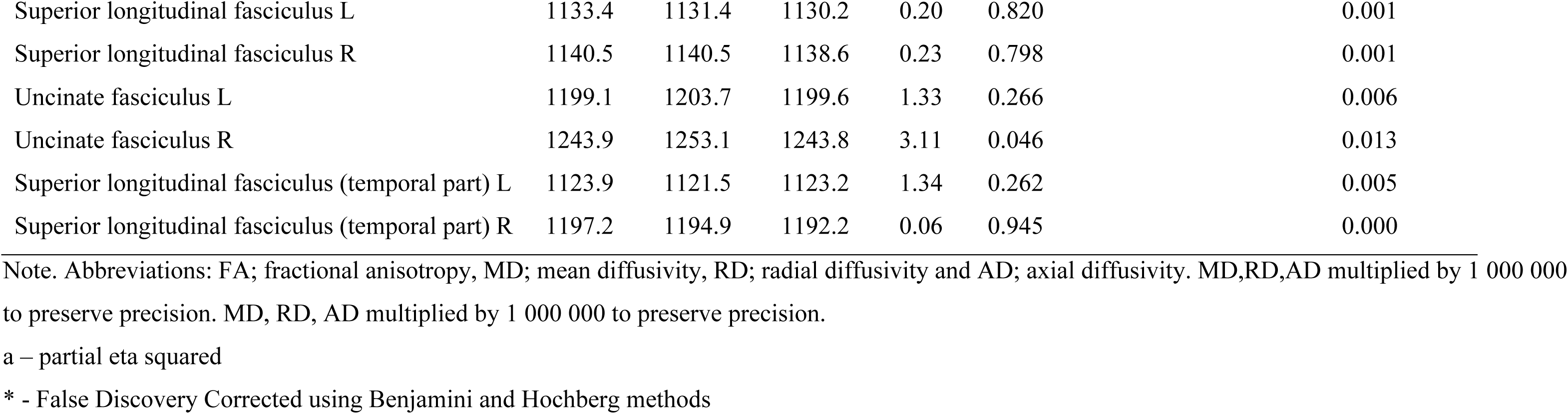
Anatomical regions of interest (ROI) analyses

| FA | SZ | BD | HC | F | *p* | Pairwise comparison | $\eta^2$ <sup>(a)</sup> |
| --- | --- | --- | --- | --- | --- | --- | --- |
| Mean FA | 0.45 | 0.45 | 0.46 | 7.02 | <0.001* | HC>SZ, BD>SZ | 0.029 |
| Genu of corpus callosum | 0.69 | 0.69 | 0.70 | 3.73 | 0.025 |  | 0.015 |
| Body of corpus callosum | 0.70 | 0.70 | 0.71 | 6.15 | 0.002 | HC>SZ | 0.025 |
| Splenium of corpus callosum | 0.79 | 0.79 | 0.79 | 7.18 | <0.001* | HC>SZ, BD>SZ | 0.029 |
| Anterior thalamic radiation L | 0.46 | 0.47 | 0.47 | 6.01 | 0.003 | HC>SZ, BD>SZ | 0.025 |
| Anterior thalamic radiation R | 0.47 | 0.47 | 0.47 | 6.99 | <0.001* | HC>SZ, BD>SZ | 0.029 |
| Corticospinal tract L | 0.59 | 0.59 | 0.59 | 3.36 | 0.035 | BD>SZ | 0.014 |
| Corticospinal tract R | 0.57 | 0.58 | 0.57 | 4.05 | 0.018 | BD>SZ | 0.017 |
| Cingulum (cingulate gyrus) L | 0.50 | 0.50 | 0.51 | 5.81 | 0.003 | HC>SZ | 0.024 |
| Cingulum (cingulate gyrus) R | 0.49 | 0.49 | 0.50 | 3.84 | 0.022 | HC>SZ | 0.016 |
| Cingulum (hippocampus) L | 0.41 | 0.42 | 0.42 | 1.72 | 0.181 |  | 0.007 |
| Cingulum (hippocampus) R | 0.43 | 0.44 | 0.44 | 1.98 | 0.139 |  | 0.008 |
| Forceps major | 0.54 | 0.54 | 0.54 | 6.98 | <0.001* | HC>SZ | 0.029 |
| Forceps minor | 0.50 | 0.50 | 0.51 | 4.46 | 0.012 | HC>SZ | 0.018 |
| Inferior fronto-occipital fasciculus L | 0.46 | 0.46 | 0.47 | 6.71 | <0.001* | HC>SZ | 0.027 |
| Inferior fronto-occipital fasciculus R | 0.47 | 0.47 | 0.48 | 7.51 | <0.001* | HC>SZ, BD>SZ | 0.031 |
| Inferior longitudinal fasciculus L | 0.43 | 0.44 | 0.44 | 6.12 | 0.002 | HC>SZ, BD>SZ | 0.025 |
| Inferior longitudinal fasciculus R | 0.40 | 0.40 | 0.41 | 7.94 | <0.001* | HC>SZ, BD>SZ | 0.032 |
| Superior longitudinal fasciculus L | 0.44 | 0.44 | 0.44 | 5.63 | 0.004 | HC>SZ | 0.023 |
| Superior longitudinal fasciculus R | 0.44 | 0.44 | 0.44 | 6.20 | 0.002 | HC>SZ, BD>SZ | 0.025 |
| Uncinate fasciculus L | 0.41 | 0.41 | 0.42 | 2.38 | 0.093 |  | 0.010 |
| Uncinate fasciculus R | 0.42 | 0.43 | 0.42 | 4.61 | 0.010 |  | 0.019 |
| Superior longitudinal fasciculus (temporal part) L | 0.44 | 0.44 | 0.45 | 4.02 | 0.019 | HC>SZ, BD>SZ | 0.017 |
| Superior longitudinal fasciculus (temporal part) R | 0.48 | 0.48 | 0.48 | 2.48 | 0.085 |  | 0.010 |
| MD |  |  |  |  |  |  |  |
| | SZ | BD | HC | F | *p* | Pairwise comparison | $\eta^2$ <sup>(a)</sup> |
| Mean MD | 782.0 | 782.1 | 776.7 | 0.83 | 0.437 |  | 0.003 |
| Genu of corpus callosum | 855.3 | 869.3 | 856.5 | 1.01 | 0.364 |  | 0.004 |
| Body of corpus callosum | 822.6 | 820.1 | 811.4 | 2.65 | 0.071 |  | 0.011 |
| Splenium of corpus callosum | 722.8 | 717.4 | 715.1 | 1.77 | 0.171 |  | 0.007 |
| Anterior thalamic radiation L | 790.6 | 793.4 | 783.5 | 0.58 | 0.560 |  | 0.002 |
| Anterior thalamic radiation R | 808.2 | 812.5 | 800.5 | 1.98 | 0.139 |  | 0.008 |
| Corticospinal tract L | 709.7 | 703.2 | 703.8 | 1.65 | 0.192 |  | 0.007 |
| Corticospinal tract R | 714.0 | 707.3 | 709.0 | 1.94 | 0.146 |  | 0.008 |
| Cingulum (cingulate gyrus) L | 767.2 | 767.3 | 762.6 | 0.50 | 0.606 |  | 0.002 |
| Cingulum (cingulate gyrus) R | 769.3 | 767.4 | 764.8 | 0.81 | 0.444 |  | 0.003 |

Tønnesen
| Cingulum (hippocampus) L | 801.6 | 801.9 | 796.8 | 0.10 | 0.904 |  | 0.000 |
| --- | --- | --- | --- | --- | --- | --- | --- |
| Cingulum (hippocampus) R | 818.3 | 815.9 | 811.6 | 0.19 | 0.824 |  | 0.001 |
| Forceps major | 768.5 | 765.0 | 760.2 | 3.61 | 0.028 |  | 0.015 |
| Forceps minor | 847.6 | 858.9 | 847.4 | 2.01 | 0.135 |  | 0.008 |
| Inferior fronto-occipital fasciculus L | 793.9 | 797.6 | 789.1 | 1.19 | 0.305 |  | 0.005 |
| Inferior fronto-occipital fasciculus R | 798.1 | 798.6 | 793.1 | 1.25 | 0.287 |  | 0.005 |
| Inferior longitudinal fasciculus L | 780.3 | 780.1 | 775.2 | 0.38 | 0.682 |  | 0.002 |
| Inferior longitudinal fasciculus R | 793.1 | 795.5 | 789.2 | 0.79 | 0.455 |  | 0.003 |
| Superior longitudinal fasciculus L | 753.7 | 752.1 | 747.7 | 1.25 | 0.286 |  | 0.005 |
| Superior longitudinal fasciculus R | 757.6 | 756.7 | 752.5 | 0.94 | 0.390 |  | 0.004 |
| Uncinate fasciculus L | 811.5 | 813.2 | 808.6 | 0.03 | 0.974 |  | 0.000 |
| Uncinate fasciculus R | 829.9 | 831.5 | 826.3 | 0.14 | 0.870 |  | 0.001 |
| Superior longitudinal fasciculus (temporal part) L | 744.4 | 742.4 | 739.8 | 0.47 | 0.623 |  | 0.002 |
| Superior longitudinal fasciculus (temporal part) R | 761.2 | 760.5 | 755.8 | 0.79 | 0.455 |  | 0.003 |
| RD |  |  |  |  |  |  |  |
| | SZ | BD | HC | F | *p* | Pairwise comparison | $\eta^2$<sup>(a)</sup> |
| Mean RD | 571.8 | 571.1 | 564.5 | 2.960 | 0.053 |  | 0.012 |
| Genu of corpus callosum | 439.3 | 449.8 | 433.0 | 1.740 | 0.177 |  | 0.007 |
| Body of corpus callosum | 417.3 | 411.8 | 401.3 | 5.023 | 0.007 | HC<SZ | 0.021 |
| Splenium of corpus callosum | 290.6 | 279.3 | 279.6 | 5.477 | 0.004 | HC<SZ, BD<SZ | 0.023 |
| Anterior thalamic radiation L | 577.5 | 578.7 | 568.0 | 1.710 | 0.182 |  | 0.007 |
| Anterior thalamic radiation R | 589.1 | 592.2 | 578.9 | 3.752 | 0.024 |  | 0.016 |
| Corticospinal tract L | 443.1 | 436.3 | 437.8 | 2.713 | 0.067 |  | 0.011 |
| Corticospinal tract R | 453.5 | 446.2 | 448.5 | 3.417 | 0.034 | BD<SZ | 0.014 |
| Cingulum (cingulate gyrus) L | 527.2 | 527.5 | 518.4 | 3.142 | 0.044 | HC<SZ | 0.013 |
| Cingulum (cingulate gyrus) R | 539.0 | 538.9 | 531.4 | 2.595 | 0.076 |  | 0.011 |
| Cingulum (hippocampus) L | 615.3 | 614.0 | 608.1 | 0.131 | 0.877 |  | 0.001 |
| Cingulum (hippocampus) R | 615.5 | 612.4 | 606.9 | 0.741 | 0.477 |  | 0.003 |
| Forceps major | 496.6 | 491.5 | 485.8 | 6.376 | 0.002 | HC<SZ | 0.026 |
| Forceps minor | 582.8 | 591.8 | 578.5 | 1.666 | 0.190 |  | 0.007 |
| Inferior fronto-occipital fasciculus L | 576.7 | 578.6 | 568.8 | 3.308 | 0.037 | HC<SZ | 0.014 |
| Inferior fronto-occipital fasciculus R | 573.5 | 572.1 | 566.0 | 3.722 | 0.025 | HC<SZ | 0.015 |
| Inferior longitudinal fasciculus L | 581.6 | 579.6 | 574.0 | 2.071 | 0.127 |  | 0.009 |
| Inferior longitudinal fasciculus R | 611.4 | 612.3 | 604.9 | 2.374 | 0.094 |  | 0.010 |
| Superior longitudinal fasciculus L | 563.9 | 562.4 | 556.4 | 2.915 | 0.055 |  | 0.012 |
| Superior longitudinal fasciculus R | 566.1 | 564.8 | 559.5 | 2.635 | 0.073 |  | 0.011 |
| Uncinate fasciculus L | 617.7 | 617.9 | 613.1 | 0.439 | 0.645 |  | 0.002 |
| Uncinate fasciculus R | 622.9 | 620.7 | 617.6 | 1.092 | 0.336 |  | 0.005 |
| Superior longitudinal fasciculus (temporal part) L | 554.7 | 552.9 | 548.1 | 1.749 | 0.175 |  | 0.007 |
| Superior longitudinal fasciculus (temporal part) R | 543.3 | 543.4 | 537.6 | 1.209 | 0.299 |  | 0.005 |

| AD |  |  |  |  |  |  |  |
| --- | --- | --- | --- | --- | --- | --- | --- |
| | SZ | BD | HC | F | *p* | Pairwise comparison | $\eta p^2$ <sup>(a)</sup> |
| Mean AD | 1202.5 | 1204.2 | 1201.2 | 0.71 | 0.492 |  | 0.003 |
| Genu of corpus callosum | 1687.3 | 1708.2 | 1703.5 | 4.81 | 0.009 |  | 0.020 |
| Body of corpus callosum | 1633.1 | 1636.7 | 1631.6 | 0.38 | 0.688 |  | 0.002 |
| Splenium of corpus callosum | 1587.2 | 1593.7 | 1586.0 | 0.72 | 0.488 |  | 0.003 |
| Anterior thalamic radiation L | 1216.8 | 1222.6 | 1214.4 | 0.27 | 0.764 |  | 0.001 |
| Anterior thalamic radiation R | 1246.4 | 1253.0 | 1243.6 | 0.67 | 0.512 |  | 0.003 |
| Corticospinal tract L | 1242.9 | 1237.0 | 1235.8 | 0.31 | 0.734 |  | 0.001 |
| Corticospinal tract R | 1235.1 | 1229.6 | 1229.9 | 0.21 | 0.808 |  | 0.001 |
| Cingulum (cingulate gyrus) L | 1247.1 | 1246.9 | 1250.9 | 2.65 | 0.072 |  | 0.011 |
| Cingulum (cingulate gyrus) R | 1229.9 | 1224.5 | 1231.4 | 0.86 | 0.425 |  | 0.004 |
| Cingulum (hippocampus) L | 1174.2 | 1177.8 | 1174.3 | 0.68 | 0.508 |  | 0.003 |
| Cingulum (hippocampus) R | 1223.7 | 1222.8 | 1221.0 | 0.12 | 0.890 |  | 0.001 |
| Forceps major | 1312.4 | 1312.1 | 1308.9 | 0.20 | 0.819 |  | 0.001 |
| Forceps minor | 1377.3 | 1393.0 | 1385.2 | 7.33 | <0.001* |  | 0.030 |
| Inferior fronto-occipital fasciculus L | 1228.3 | 1235.5 | 1229.6 | 2.33 | 0.099 |  | 0.010 |
| Inferior fronto-occipital fasciculus R | 1247.3 | 1251.7 | 1247.1 | 1.08 | 0.340 |  | 0.005 |
| Inferior longitudinal fasciculus L | 1177.6 | 1181.2 | 1177.6 | 0.78 | 0.458 |  | 0.003 |
| Inferior longitudinal fasciculus R | 1156.3 | 1162.0 | 1157.9 | 2.10 | 0.124 |  | 0.009 |

Tønnesen
|  |  |  |  |  |  |  |
| --- | --- | --- | --- | --- | --- | --- |
| Superior longitudinal fasciculus L | 1133.4 | 1131.4 | 1130.2 | 0.20 | 0.820 | 0.001 |
| Superior longitudinal fasciculus R | 1140.5 | 1140.5 | 1138.6 | 0.23 | 0.798 | 0.001 |
| Uncinate fasciculus L | 1199.1 | 1203.7 | 1199.6 | 1.33 | 0.266 | 0.006 |
| Uncinate fasciculus R | 1243.9 | 1253.1 | 1243.8 | 3.11 | 0.046 | 0.013 |
| Superior longitudinal fasciculus (temporal part) L | 1123.9 | 1121.5 | 1123.2 | 1.34 | 0.262 | 0.005 |
| Superior longitudinal fasciculus (temporal part) R | 1197.2 | 1194.9 | 1192.2 | 0.06 | 0.945 | 0.000 |

Note. Abbreviations: FA; fractional anisotropy, MD; mean diffusivity, RD; radial diffusivity and AD; axial diffusivity. MD,RD,AD multiplied by 1 000 000 to preserve precision. MD, RD, AD multiplied by 1 000 000 to preserve precision.
a – partial eta squared
\* - False Discovery Corrected using Benjamini and Hochberg methods

**Figure 4.**
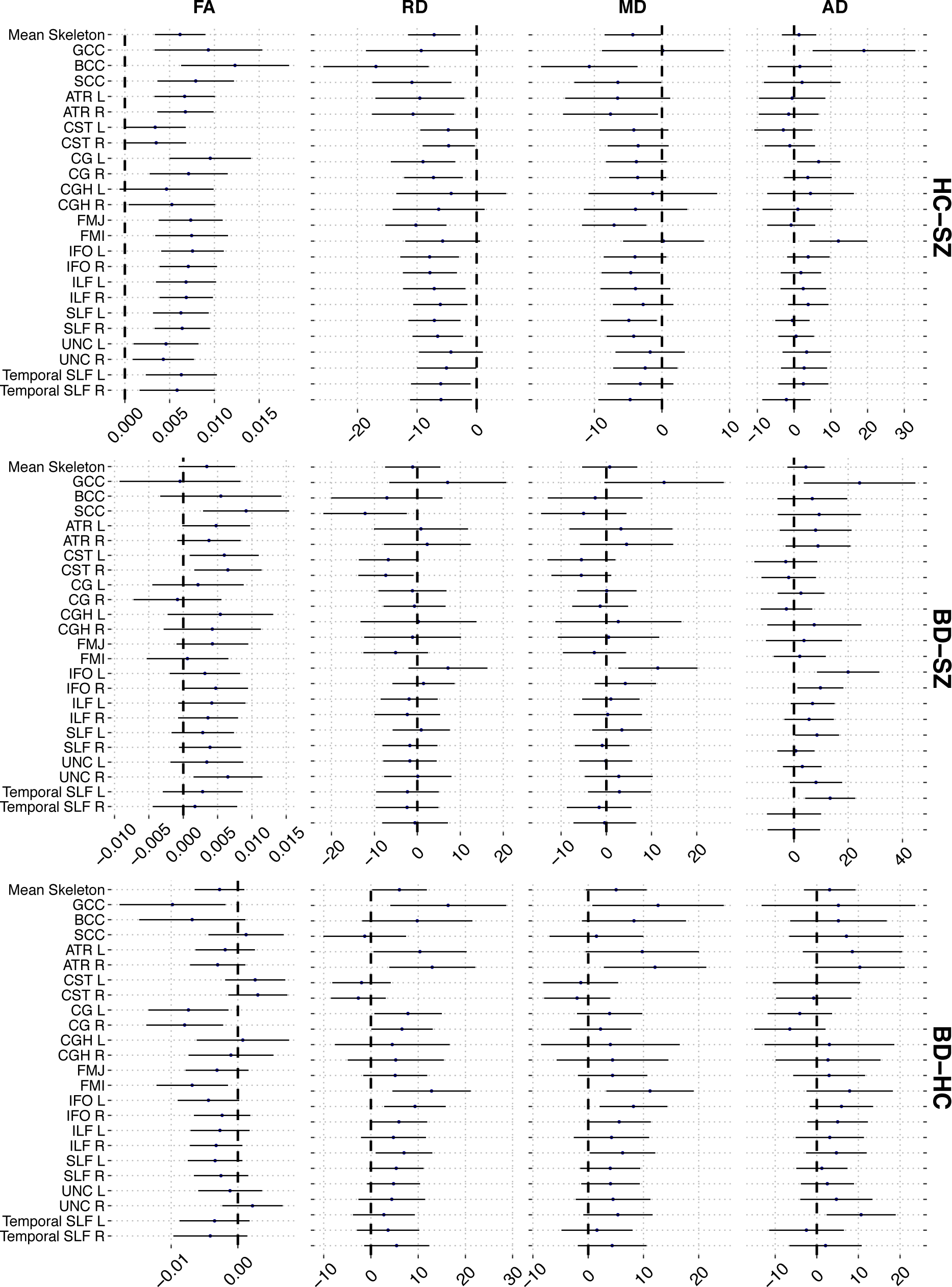
Results from region of interest (ROI) analyses with mean difference and variance from pairwise comparisons plotted for each DTI metric. The error bars represent 95% confidence intervals.

List of abbreviations: Genu of corpus callosum (GCC), Body of corpus callosum (BCC), Splenium of corpus callosum (SCC), Anterior thalamic radiation L (ATR L), Anterior thalamic radiation R (ATR R), Corticospinal tract L (CST L), Corticospinal tract R (CST R), Cingulum (cingulate gyrus) L (CGL), Cingulum (cingulate gyrus) R (CG R), Cingulum (hippocampus) L (CGH L), Cingulum (hippocampus) R (CGH R), Forceps major (FMJ), Forceps minor (FMI), Inferior fronto-occipital fasciculus L (IFO L), Inferior fronto-occipital fasciculus R (IFO R), Inferior longitudinal fasciculus L (ILF L), Inferior longitudinal fasciculus R (ILF R), Superior longitudinal fasciculus L (SLF L), Superior longitudinal fasciculus R (SLF R), Uncinate fasciculus L (UNC L), Uncinate fasciculus R (UNC R), Superior longitudinal fasciculus (temporal part) L (Temporal SLF L), Superior longitudinal fasciculus (temporal part) R(Temporal SLF R).

## Discussion

One of the major implications of the brain dysconnectivity hypothesis of psychotic disorders is that the WM microstructural layout and integrity modulates risk and give rise to a range of symptoms. In order to test this hypothesis, we compared DTI metrics between patients with SZ and HC across the brain. Our inclusion of a group of patients with BD allowed us to test for diagnostic specificity or, conversely, cross-diagnostic convergence. In line with our primary hypothesis and converging evidence^16^, the results revealed robust differences between patients with SZ and HC on several metrics, in particular lower FA across the brain in patients, even after careful quality assessment. In general, the results were not strongly dependent on age or sex and we found no significant associations with symptoms across groups. Adding to the accumulated evidence of brain gray matter abnormalities in patients with severe mental illness^52,59^, these results support converging evidence implicating WM abnormalities in SZ and suggest these abnormalities are more pronounced for SZ than for BD.

Clinical overlaps have motivated a dimensional approach to reveal common and distinct disease mechanisms in SZ and BD. In line with cognitive and genetic studies^23,24^ suggesting several commonalities, brain imaging has not revealed structural or functional brain characteristics unambiguously distinguishing the two disorders, and previous DTI studies comparing SZ and BD have been largely inconclusive^32,34,35,60^.

The neurobiological underpinnings of DTI metrics are complex and multidimensional, and our findings do not allow for interpretation regarding specific cellular processes. Previous studies have shown associations between RD and myelin related processes^61,62^, and higher RD may suggest reduced myelin integrity in SZ, in particular when considered in light of genetic studies reporting altered expression of genes involved in lipid homeostasis and myelination^9,63^. Complementary models implicate microglial inflammatory processes and oxidative stress in WM pathology^64,65^, and inflammation-related cytokines and growth factors have been associated with reduced FA and increased RD and MD in BD ^66^. Further studies are needed to delineate the roles of myelination and inflammation for WM integrity and mental health across the lifespan^67,68^.

Considering the strong impact of age on brain WM microstructure^69,70^, characterizing age trajectories within groups may provide indirect information about the temporal evolution of aberrations in an ontogenetic perspective. Both neurodevelopmental and neurodegenerative models of the development and sustainment of psychosis have been formulated^46,47,71^, and both genetic liability and neurodevelopmental perturbations play critical roles in the modulation of risk^43,44^. Indeed, the emergence of psychotic symptoms in late adolescence and early adulthood may in fact reflect late stages of the disorder^42^. Patients with early onset SZ show microstructural aberrations^72^, which could manifest as delayed WM development during adolescence^48^. The current lack of group-by-age interactions may suggest the observed group differences are explained by events prior to the sampled age-range, which would indicate parallel age trajectories in adulthood^47^.

For HC, we observed a peak of FA at approx. 32 years followed by decreases until the maximal sampled age, which is highly corresponding with previous cross-sectional studies^69^. The FA trajectory for SZ (approx. 27 years) and BD (approx. 27 years) showed an earlier peak, followed by a linear decrease until the maximum age. Although permutation testing revealed no significant differences in age at peak FA, the sliding window approach, which provide further insight beyond a standard age-by-diagnosis interaction test, suggested the magnitude of the group differences are not completely invariant to age, with indications of increasing group differences in FA between SZ and HC until the late 20s. In addition to suggesting early and possibly accelerating age-related differences in SZ, moderate age-related differences in effect sizes may also reflect a combination of clinical heterogeneity, sampling issues, and power. The lack of significant age by group interactions possibly also hints at the shortcomings of simple models for delineating and comparing complex trajectories^73^. Future studies utilizing a longitudinal design including a wide age-range and participants at genetic or clinical risk who have still not developed psychosis are needed to characterize the trajectories of the dynamic WM aberrations during the course of brain maturation and disease development^22^.

Although detrimental effects of subject motion and other sources of noise on DTI metrics have been documented^53,55,74^, most previous DTI studies on psychotic disorders have not provided sufficient details regarding the employed QC procedures or included any quantitative QC measures as covariates. It is therefore largely unknown to which degree various sources of noise have contributed to the reported group differences. A major strength of this study is the use of an automated approach for identifying and replacing slices with signal loss due to bulk motion, considerably increasing temporal signal-to-noise ratio (tSNR^21^), and a comprehensive QC protocol including both manual and automated quantitative measures. Comparisons of summary statistics and group differences at different steps in the QC and exclusion procedure revealed a tendency of increasing group differences in FA between SZ and HC with the exclusion of noisy data. These results indicate that stringent QC may increase sensitivity to WM aberrations in SZ, and suggest that future studies should carefully address different sources of noise in their datasets before interpreting their findings as reflecting relevant pathophysiology.

Some limitations should be considered while interpreting our findings. The influence of medication on WM is debated^75-77^. As most patients were medicated, with the majority of patients taking antipsychotics, confounding effects of medication cannot be ruled out. Future studies with an appropriate design for assessing medication effects are needed. Despite our current lack of significant sex by diagnosis interaction, there is a growing appreciation of sexual dimorphisms in brain and behavior both in health and disease, which warrants further investigations. Further, our cross-sectional design is not suitable for delineating dynamic individual changes in WM microstructure, and further studies utilizing a prospective design in younger children and adolescents are needed to map microstructural changes to risk and development of psychosis. Integrating a wider range of MRI modalities with clinical, cognitive and genetic features^21,78^, and including microstructural indices based on multi-compartment diffusion models (e.g., Neurite Orientation Dispersion and Density Imaging^79,80^, free water imaging ^81^, or restriction spectrum imaging^82^), cortical and subcortical morphometry and functional measures, may prove helpful for increasing diagnostic sensitivity and specificity.

In conclusion, we report widespread WM microstructural aberration in patients with SZ compared to BD and HC. We found no significant differences between patients with BD and HC, suggesting the biophysical processes causing DTI based WM abnormalities in severe mental disorders are more prominent for SZ. These results are in line with converging genetic and pathological evidence implicating neuroinflammatory and lipid and myelin processes in SZ pathophysiology.

## Methods

### Sample

Adult patients were recruited from psychiatric units in four major hospitals Oslo. Patients had to fulfill criteria for a Structured Clinical Interview (SCID) ^83^ DSM-IV diagnosis of schizophrenia spectrum disorder, collectively referred to as SZ (n=128 including schizophrenia (n=70), schizoaffective (n=18), schizophreniform (n=7)) and psychosis not otherwise specified (n=33), or bipolar spectrum disorder, collectively referred to as BD (n=61 including BDI (n=39), BDII (n=17) and BD NOS (n=5)). The sample comprised both medicated (n=133), unmedicated (n=7) and patients missing information regarding medication status (n=49).

293 healthy controls from the same catchment area were invited through a stratified randomized selection from the national records. Exclusion criteria for both patients and HC included hospitalized head trauma, neurological disorder or IQ below 70. In addition, HC were screened with a questionnaire about severe mental illness and the Primary Care Evaluation of Mental Disorders (PRIME-MD)^84^. Exclusion criteria included somatic disease, substance abuse or dependency the last 12 months or a first-degree relative with a lifetime history of severe psychiatric disorder (SZ, BD, or major depressive disorder). The Tematisk Område Psykoser (TOP) Study is approved by the Regional Ethics Committtee (REK Sør-Øst C, 2009/2485) and the Norwegian Data Inspectorate (2003/2052). Study protocol and procedures adhered to the ethics approval and to the Declaration of Helsinki. All participants provided written informed consent, see SI for more information regarding neuropsychological and clinical assessment. Due to confidentiality and privacy of participant information data may not be shared readily online, but data can be requested by contacting the authors.

### MRI acquisition

Imaging was performed on a General Electric (Signa HDxt) 3T scanner using an 8-channel head coil at Oslo University Hospital. For DTI, a 2D spin-echo whole-brain echo planar imaging pulse with the following parameters was used: repetition time: 15 s; echo time: 85ms; flip angle: 90°; slice thickness: 2.5 mm; in-plane resolution: 1.875*1.875 mm; 30 volumes with different gradient directions (b=1000 s/mm^2^) in addition to two b=0 volumes with reversed phase-encode (blip up/down) were acquired.

### DTI processing

Image analyses and tensor calculations were performed using FSL^85-87^. Pre-processing steps included topup (http://fsl.fmrib.ox.ac.uk/fsl/fslwiki/topup)^88^ and eddy (http://fsl.fmrib.ox.ac.uk/fsl/fslwiki/eddy)^89,90^ to correct for geometrical distortions and eddy currents. Topup uses information from the reversed phase-encode blips, resulting in pairs of images with distortions going in opposite directions. From these image pairs the susceptibility-induced off-resonance field was estimated and the two images were combined into a single corrected one. Eddy detects and replaces slices affected by signal loss due to bulk motion during diffusion encoding, which is performed within an integrated framework along with correction for susceptibility induced distortions, eddy currents and motion^90^. In order to assess the effect of replacement of dropout-slices on tSNR we also processed the data using eddy without slice replacement (Supplementary Fig. S11). Briefly, mean tSNR was significantly (t=25.76, p<.001) lower when running eddy without slice replacement (mean: 7.77, SD: 0.52) compared to with slice replacement (mean 8.79, SD: 0.70). There was no significant group differences in the amount of slices replaced (F=1.046, p=0.352, mean group slice replacement: HC: 10.92 (± 7.36), BD: 10.52 (± 7.40), SZ: 12.46 (± 9.42)).

Diffusion tensor fitting was done using dtifit in FSL. FA is a scalar value of diffusion directionality, MD was computed as the mean of all three eigenvalues, RD as the mean of the second and third eigenvalue^91^, while AD represent the principal eigenvalue.

Prior to statistical analyses we employed a stepwise QC procedure, including maximum voxel intensity outlier count (MAXVOX)^92^ and tSNR^92^. Since reduced data quality due to subject motion and other factors may bias the results in clinical studies, we defined various quantitative QC metrics and tested for group differences within different QC strata. Specifically, we devised a semi-qualitative QC protocol including methods provided in DTIPrep^93^ and tSNR^94^. Supplementary Fig. S12 shows a flowchart of the QC protocol. At each step the distributions of the quality metrics were visually inspected. In our step-wise exclusion protocol, datasets were excluded based on a summary score utilizing (1) maximum MAXVOX^92^ and (2) tSNR^92^. The summary score was formed by first inverting the MAXVOX score, z-normalize both scores independently, add 10 to each of the z-scores (to avoid negative values), and then computing the product of the two. This product was then z-normalised, with low scores indicating worse quality. In an iterative fashion, subjects with a QC sum z-score below -2.5 were excluded, and the group statistics were recomputed. This was repeated until no datasets had a z-score below -2.5. Briefly, the slice-wise check and the MAXVOX screens the DWI data for intensity related artifacts while tSNR is a global summary measure. See SI for further details regarding the QC such as summary stats for each step of the QC procedure (Supplementary Table S5), demographic overview of excluded participants (Supplementary Table S6), density plots of DTI metrics before and after exclusion (Supplementary Fig. S13) and voxel-wise analyses after QC (Supplementary Fig.S9). In short, a thorough inspection of the excluded and included participants after QC suggested that general quality of the data is good. Thus, we present results on the full dataset with supplemental and complementary results from a stringent QC.

Voxelwise analysis of FA, MD, AD and RD were carried out using TBSS^51^. FA volumes were skull-stripped and aligned to the FMRIB58_FA template supplied by FSL using nonlinear registration (FNIRT)^95^. Next, mean FA were derived and thinned to create a mean FA skeleton, representing the center of all tracts common across subjects. The same warping and skeletonization was repeated for MD, AD and RD. We thresholded and binarized the mean FA skeleton at FA>0.2 before feeding the data into voxelwise statistics.

### Statistical analyses

Voxelwise statistical analyses were performed using permutation testing, implemented in FSL’s randomise^96^. Main effects of diagnosis on FA, RD, MD and AD were tested using general linear models (GLM) by forming pairwise group contrasts and corresponding F-tests. Since previous studies have documented strong curvilinear relationships between DTI features and age throughout the adult lifespan^69^, we included age, age^2^ and sex as covariates. The data was tested against an empirical null distribution generated by 5000 permutations and threshold free cluster enhancement (TFCE)^97^ was used to avoid arbitrarily defining the cluster-forming threshold. Voxelwise maps were thresholded at *p*<.05 and corrected for multiple comparisons across space. Mean FA, MD, RD and AD across the brain and within significant clusters were submitted to R^98^ for peak estimation and to compute effect sizes and visualization. In a resampling with replacement (bootstrapping) procedure we fitted the DTI data to age using local polynomial regression function (LOESS). LOESS has previously been used in lifespan studies^69^ and avoids some of the shortcoming of polynomial models for age fitting^73^. Using boot package^99,100^ in R, we repeated the age fitting procedure for each of the 10,000 bootstrapped samples for each group to estimate the mean age at the maximum (FA) or minimum (MD, RD, AD) value across iterations, and its uncertainty with confidence intervals calculated using the adjusted bootstrap percentile method. Additionally, the group differences in age at peak FA was tested against an empirical null distribution generated by 10000 permutations, generated by randomly shuffling group labels and computing the pairwise group differences at each iteration. All pairwise differences were combined into one null distribution, and the differences in the true data were compared to this common null, enabling correction for multiple comparisons across all pairwise comparisons.

We tested for associations between GAF/PANSS domains and FA, MD, RD and AD across both patient groups (SZ and BD grouped together) in the whole brain and within specific regions, covarying for age, age^2^ and sex. False discovery rate (FDR)^101^ and Bonferroni was used to correct for multiple testing.

Differences between and within subgroups (see SI for more information), group by age and group by age^2^ interactions on the mean skeleton DTI metrics were tested. In order to account for heterogeneity in the diagnostic groups we ran subgroup analyses on the mean skeleton metrics on the largest subgroups (strict SZ, PNOS, BDI and BDII). Additionally, in a control analysis to confirm that possible differences in age distribution did not influence the main results we excluded participants over 55 years of age and ran mean skeleton analyses across groups.

We performed a sliding window technique to obtain effect sizes for each of the pairwise group comparison within different age-strata. Utilizing the zoo R package^102^, we slid a window of 150 participants in steps of 5 participants along the sorted age span. At each step, we computed a linear model investigating effects of diagnosis, accounting for sex. We plotted the resulting t-values and effect sizes (Cohen’s d) representing pairwise group differences against the mean age of each sliding group and fit a LOESS function using ggplot2 in R^103^. In order to test if group differences varied between females and males we reran the analysis when including a sex-by-diagnosis interaction term for the mean skeleton and ROI analyses.

To facilitate future meta-analyses, we calculated raw mean DTI values across the skeleton and within various anatomical regions of interest (ROIs) based on the intersection between the TBSS skeleton and probabilistic atlases^104,105^. R was used for further analysis, including linear models with each of the ROI DTI value as dependent variable, diagnostic group and sex as fixed factors, and age and age^2^ as covariates.

## Acknowledgements

This work was funded by the South-Eastern Norway Regional Health Authority (2014097, 2015073, 2016083), the Research Council of Norway (204966, 249795, 223273), KG Jebsen Stiftelsen and the European Commission’s 7th Framework Programme (#602450, IMAGEMEND).

## Author Contributions Statement

ST and LTW designed the study. ST, TK, and LTW performed the analyses with contributions from DA and NTD. ST, LTW and TK interpreted the results. ST, LTW and TK drafted the manuscript. All authors reviewed and approved the manuscript

## References

1 Vos, T. et al. Years lived with disability (YLDs) for 1160 sequelae of 289 diseases and injuries 1990-2010: a systematic analysis for the Global Burden of Disease Study 2010. Lancet 380, 2163–2196, doi:10.1016/S0140-6736(12)61729-2 (2013).

2 Coyle, J. T., Balu, D. T., Puhl, M. D. & Konopaske, G. T. History of the Concept of Disconnectivity in Schizophrenia. Harvard review of psychiatry 24, 80–86, doi:10.1097/hrp.0000000000000102 (2016).

3 Friston, K. J. The disconnection hypothesis. Schizophrenia research 30, 115–125 (1998).

4 Stephan, K. E., Friston, K. J. & Frith, C. D. Dysconnection in schizophrenia: from abnormal synaptic plasticity to failures of self-monitoring. Schizophrenia bulletin 35, 509–527, doi:10.1093/schbul/sbn176 (2009).

5 Kaufmann, T. et al. Disintegration of sensorimotor brain networks in schizophrenia. Schizophrenia bulletin, doi:10.1093/schbul/sbv060 (2015).

6 Tkachev, D. et al. Oligodendrocyte dysfunction in schizophrenia and bipolar disorder. Lancet 362, 798–805, doi:10.1016/S0140-6736(03)14289-4 (2003).

7 Hakak, Y. et al. Genome-wide expression analysis reveals dysregulation of myelination-related genes in chronic schizophrenia. Proceedings of the National Academy of Sciences of the United States of America 98, 4746–4751, doi:10.1073/pnas.081071198 (2001).

8 Konrad, A. & Winterer, G. Disturbed structural connectivity in schizophrenia primary factor in pathology or epiphenomenon? Schizophrenia bulletin 34, 72–92, doi:10.1093/schbul/sbm034 (2008).

9 Steen, V. M. et al. Genetic evidence for a role of the SREBP transcription system and lipid biosynthesis in schizophrenia and antipsychotic treatment. European neuropsychopharmacology: the journal of the European College of Neuropsychopharmacology, doi:10.1016/j.euroneuro.2016.07.011 (2016).

10 Stedehouder, J. & Kushner, S. A. Myelination of parvalbumin interneurons: a parsimonious locus of pathophysiological convergence in schizophrenia. Molecular psychiatry, doi:10.1038/mp.2016.147 (2016).

11 van Kesteren, C. F. et al. Immune involvement in the pathogenesis of schizophrenia: a meta-analysis on postmortem brain studies. Translational psychiatry 7, e1075, doi:10.1038/tp.2017.4 (2017).

12 Klauser, P. et al. White Matter Disruptions in Schizophrenia Are Spatially Widespread and Topologically Converge on Brain Network Hubs. Schizophrenia bulletin, doi:10.1093/schbul/sbw100 (2016).

13 Ellison-Wright, I. & Bullmore, E. Meta-analysis of diffusion tensor imaging studies in schizophrenia. Schizophrenia research 108, 3–10, doi:10.1016/j.schres.2008.11.021 (2009).

14 Canu, E., Agosta, F. & Filippi, M. A selective review of structural connectivity abnormalities of schizophrenic patients at different stages of the disease. Schizophrenia research 161, 19–28, doi:10.1016/j.schres.2014.05.020 (2015).

15 Patel, S. et al. A meta-analysis of diffusion tensor imaging studies of the corpus callosum in schizophrenia. Schizophrenia research 129, 149–155, doi:10.1016/j.schres.2011.03.014 (2011).

16 Kelly, S. et al. Widespread white matter microstructural differences in schizophrenia across 4322 individuals: results from the ENIGMA Schizophrenia DTI Working Group. Molecular psychiatry, doi:10.1038/mp.2017.170 (2017).

17 Moon, C. M. & Jeong, G. W. Abnormalities in gray and white matter volumes associated with explicit memory dysfunction in patients with generalized anxiety disorder. Acta radiologica (Stockholm, Sweden: 1987) 58, 353–361, doi:10.1177/0284185116649796 (2017).

18 Gan, J. et al. Abnormal white matter structural connectivity in adults with obsessive-compulsive disorder. Translational psychiatry 7, e1062, doi:10.1038/tp.2017.22 (2017).

19 Jiang, J. et al. Microstructural brain abnormalities in medication-free patients with major depressive disorder: a systematic review and meta-analysis of diffusion tensor imaging. Journal of psychiatry & neuroscience: JPN 42, 150341, doi:10.1503/jpn.150341 (2016).

20 Westlye, L. T., Bjornebekk, A., Grydeland, H., Fjell, A. M. & Walhovd, K. B. Linking an anxiety-related personality trait to brain white matter microstructure: diffusion tensor imaging and harm avoidance. Archives of general psychiatry 68, 369–377, doi:10.1001/archgenpsychiatry.2011.24 (2011).

21 Doan, N. T. et al. Dissociable diffusion MRI patterns of white matter microstructure and connectivity in Alzheimer’s disease spectrum. Scientific reports 7, 45131, doi:10.1038/srep45131 (2017).

22 Alnæs, D. et al. Association of heritable cognitive ability and psychopathology with white matter properties in children and adolescents. JAMA psychiatry, doi:10.1001/jamapsychiatry.2017.4277 (2018).

23 Owen, M. J., Craddock, N. & Jablensky, A. The genetic deconstruction of psychosis. Schizophrenia bulletin 33, 905–911, doi:10.1093/schbul/sbm053 (2007).

24 Hill, S. K. et al. A comparison of neuropsychological dysfunction in first-episode psychosis patients with unipolar depression, bipolar disorder, and schizophrenia. Schizophrenia research 113, 167–175, doi:10.1016/j.schres.2009.04.020 (2009).

25 Simonsen, C. et al. Neurocognitive dysfunction in bipolar and schizophrenia spectrum disorders depends on history of psychosis rather than diagnostic group. Schizophrenia bulletin 37, 73–83, doi:10.1093/schbul/sbp034 (2011).

26 Goldberg, T. E. Some fairly obvious distinctions between schizophrenia and bipolar disorder. Schizophrenia research 39, 127-132; discussion 161-122 (1999).

27 Lagopoulos, J. et al. Microstructural white matter changes in the corpus callosum of young people with Bipolar Disorder: a diffusion tensor imaging study. PloS one 8, e59108, doi:10.1371/journal.pone.0059108 (2013).

28 Sarrazin, S. et al. A multicenter tractography study of deep white matter tracts in bipolar I disorder: psychotic features and interhemispheric disconnectivity. JAMA psychiatry 71, 388–396, doi:10.1001/jamapsychiatry.2013.4513 (2014).

29 Mahon, K., Burdick, K. E., Wu, J., Ardekani, B. A. & Szeszko, P. R. Relationship between suicidality and impulsivity in bipolar I disorder: a diffusion tensor imaging study. Bipolar disorders 14, 80–89, doi:10.1111/j.1399-5618.2012.00984.x (2012).

30 Chen, Z. et al. Voxel based morphometric and diffusion tensor imaging analysis in male bipolar patients with first-episode mania. Progress in neuro-psychopharmacology & biological psychiatry 36, 231–238, doi:10.1016/j.pnpbp.2011.11.002 (2012).

31 Sprooten, E. et al. A comprehensive tractography study of patients with bipolar disorder and their unaffected siblings. Human brain mapping 37, 3474–3485, doi:10.1002/hbm.23253 (2016).

32 Li, J. et al. A comparative diffusion tensor imaging study of corpus callosum subregion integrity in bipolar disorder and schizophrenia. Psychiatry research 221, 58–62, doi:10.1016/j.pscychresns.2013.10.007 (2014).

33 Cui, L. et al. Assessment of white matter abnormalities in paranoid schizophrenia and bipolar mania patients. Psychiatry research 194, 347–353, doi:10.1016/j.pscychresns.2011.03.010 (2011).

34 Sussmann, J. E. et al. White matter abnormalities in bipolar disorder and schizophrenia detected using diffusion tensor magnetic resonance imaging. Bipolar disorders 11, 11–18, doi:10.1111/j.1399-5618.2008.00646.x (2009).

35 Kumar, J. et al. Shared white-matter dysconnectivity in schizophrenia and bipolar disorder with psychosis. Psychological medicine, 1-12, doi:10.1017/s0033291714001810 (2014).

36 McIntosh, A. M. et al. White matter tractography in bipolar disorder and schizophrenia. Biological psychiatry 64, 1088–1092, doi:10.1016/j.biopsych.2008.07.026 (2008).

37 Skudlarski, P. et al. Diffusion tensor imaging white matter endophenotypes in patients with schizophrenia or psychotic bipolar disorder and their relatives. The American journal of psychiatry 170, 886–898, doi:10.1176/appi.ajp.2013.12111448 (2013).

38 Mallas, E. J. et al. Genome-wide discovered psychosis-risk gene ZNF804A impacts on white matter microstructure in health, schizophrenia and bipolar disorder. PeerJ 4, e1570, doi:10.7717/peerj.1570 (2016).

39 Dong, D. et al. Shared abnormality of white matter integrity in schizophrenia and bipolar disorder: A comparative voxel-based meta-analysis. Schizophrenia research, doi:10.1016/j.schres.2017.01.005 (2017).

40 Fatemi, S. H. & Folsom, T. D. The neurodevelopmental hypothesis of schizophrenia, revisited. Schizophrenia bulletin 35, 528–548, doi:10.1093/schbul/sbn187 (2009).

41 Roybal, D. J. et al. Biological evidence for a neurodevelopmental model of pediatric bipolar disorder. The Israel journal of psychiatry and related sciences 49, 28–43 (2012).

42 Insel, T. R. Rethinking schizophrenia. Nature 468, 187–193, doi:10.1038/nature09552 (2010).

43 Paus, T., Keshavan, M. & Giedd, J. N. Why do many psychiatric disorders emerge during adolescence? Nature reviews. Neuroscience 9, 947–957, doi:10.1038/nrn2513 (2008).

44 Carletti, F. et al. Alterations in white matter evident before the onset of psychosis. Schizophrenia bulletin 38, 1170–1179, doi:10.1093/schbul/sbs053 (2012).

45 Kochunov, P. et al. Heterochronicity of white matter development and aging explains regional patient control differences in schizophrenia. Human brain mapping, doi:10.1002/hbm.23336 (2016).

46 Schnack, H. G. et al. Accelerated Brain Aging in Schizophrenia: A Longitudinal Pattern Recognition Study. The American journal of psychiatry 173, 607–616, doi:10.1176/appi.ajp.2015.15070922 (2016).

47 Kochunov, P. & Hong, L. E. Neurodevelopmental and neurodegenerative models of schizophrenia: white matter at the center stage. Schizophrenia bulletin 40, 721–728, doi:10.1093/schbul/sbu070 (2014).

48 Douaud, G. et al. Schizophrenia delays and alters maturation of the brain in adolescence. Brain: a journal of neurology 132, 2437–2448, doi:10.1093/brain/awp126 (2009).

49 Kochunov, P. et al. Testing the Hypothesis of Accelerated Cerebral White Matter Aging in Schizophrenia and Major Depression. Biological psychiatry, doi:10.1016/j.biopsych.2012.10.002 (2012).

50 Shahab, S. et al. Sex and Diffusion Tensor Imaging of White Matter in Schizophrenia: A Systematic Review Plus Meta-analysis of the Corpus Callosum. Schizophrenia bulletin, doi:10.1093/schbul/sbx049 (2017).

51 Smith, S. M. et al. Tract-based spatial statistics: voxelwise analysis of multi-subject diffusion data. NeuroImage 31, 1487–1505, doi:10.1016/j.neuroimage.2006.02.024 (2006).

52 Moberget, T. et al. Cerebellar volume and cerebellocerebral structural covariance in schizophrenia: a multisite mega-analysis of 983 patients and 1349 healthy controls. Molecular psychiatry, doi:10.1038/mp.2017.106 (2017).

53 Yendiki, A., Koldewyn, K., Kakunoori, S., Kanwisher, N. & Fischl, B. Spurious group differences due to head motion in a diffusion MRI study. NeuroImage 88, 79–90, doi:10.1016/j.neuroimage.2013.11.027 (2014).

54 Ling, J. et al. Head injury or head motion? Assessment and quantification of motion artifacts in diffusion tensor imaging studies. Human brain mapping 33, 50–62, doi:10.1002/hbm.21192 (2012).

55 Muller, H. P. et al. Impact of the control for corrupted diffusion tensor imaging data in comparisons at the group level: an application in Huntington disease. Biomedical engineering online 13, 128, doi:10.1186/1475-925x-13-128 (2014).

56 Pedersen, G., Hagtvet, K. A. & Karterud, S. Generalizability studies of the Global Assessment of Functioning-Split version. Comprehensive psychiatry 48, 88–94, doi:10.1016/j.comppsych.2006.03.008 (2007).

57 Langeveld, J. et al. Is there an optimal factor structure of the Positive and Negative Syndrome Scale in patients with first-episode psychosis? Scandinavian journal of psychology 54, 160–165, doi:10.1111/sjop.12017 (2013).

58 Wallwork, R. S., Fortgang, R., Hashimoto, R., Weinberger, D. R. & Dickinson, D. Searching for a consensus five-factor model of the Positive and Negative Syndrome Scale for schizophrenia. Schizophrenia research 137, 246–250, doi:10.1016/j.schres.2012.01.031 (2012).

59 van Erp, T. G. et al. Subcortical brain volume abnormalities in 2028 individuals with schizophrenia and 2540 healthy controls via the ENIGMA consortium. Molecular psychiatry 21, 585, doi:10.1038/mp.2015.118 (2016).

60 Lu, L. H., Zhou, X. J., Keedy, S. K., Reilly, J. L. & Sweeney, J. A. White matter microstructure in untreated first episode bipolar disorder with psychosis: comparison with schizophrenia. Bipolar disorders 13, 604–613, doi:10.1111/j.1399-5618.2011.00958.x (2011).

61 Beaulieu, C. The basis of anisotropic water diffusion in the nervous system - a technical review. NMR in biomedicine 15, 435–455, doi:10.1002/nbm.782 (2002).

62 Song, S. K. et al. Dysmyelination revealed through MRI as increased radial (but unchanged axial) diffusion of water. NeuroImage 17, 1429–1436 (2002).

63 Chavarria-Siles, I. et al. Myelination-related genes are associated with decreased white matter integrity in schizophrenia. European journal of human genetics: EJHG 24, 381–386, doi:10.1038/ejhg.2015.120 (2016).

64 Naaldijk, Y. M., Bittencourt, M. C., Sack, U. & Ulrich, H. Kinins and microglial responses in bipolar disorder: a neuroinflammation hypothesis. Biological chemistry 397, 283–296, doi:10.1515/hsz-2015-0257 (2016).

65 Najjar, S. & Pearlman, D. M. Neuroinflammation and white matter pathology in schizophrenia: systematic review. Schizophrenia research 161, 102–112, doi:10.1016/j.schres.2014.04.041 (2015).

66 Benedetti, F. et al. Inflammatory cytokines influence measures of white matter integrity in Bipolar Disorder. Journal of affective disorders 202, 1–9, doi:10.1016/j.jad.2016.05.047 (2016).

67 Pasternak, O., Kubicki, M. & Shenton, M. E. In vivo imaging of neuroinflammation in schizophrenia. Schizophrenia research 173, 200–212, doi:10.1016/j.schres.2015.05.034 (2016).

68 Pasternak, O. et al. Excessive extracellular volume reveals a neurodegenerative pattern in schizophrenia onset. The Journal of neuroscience: the official journal of the Society for Neuroscience 32, 17365–17372, doi:10.1523/jneurosci.2904-12.2012 (2012).

69 Westlye, L. T. et al. Life-span changes of the human brain white matter: diffusion tensor imaging (DTI) and volumetry. Cerebral cortex 20, 2055–2068, doi:10.1093/cercor/bhp280 (2010).

70 Sexton, C. E. et al. Accelerated changes in white matter microstructure during aging: a longitudinal diffusion tensor imaging study. The Journal of neuroscience: the official journal of the Society for Neuroscience 34, 15425–15436, doi:10.1523/JNEUROSCI.0203-14.2014 (2014).

71 Kobayashi, H. et al. Linking the developmental and degenerative theories of schizophrenia: association between infant development and adult cognitive decline. Schizophrenia bulletin 40, 1319–1327, doi:10.1093/schbul/sbu010 (2014).

72 Tamnes, C. K. & Agartz, I. White Matter Microstructure in Early-Onset Schizophrenia: A Systematic Review of Diffusion Tensor Imaging Studies. Journal of the American Academy of Child and Adolescent Psychiatry 55, 269–279, doi:10.1016/j.jaac.2016.01.004 (2016).

73 Fjell, A. M. et al. When does brain aging accelerate? Dangers of quadratic fits in cross-sectional studies. NeuroImage 50, 1376–1383, doi:10.1016/j.neuroimage.2010.01.061 (2010).

74 Haarman, B. C. et al. Diffusion tensor imaging in euthymic bipolar disorder - A tract-based spatial statistics study. Journal of affective disorders 203, 281–291, doi:10.1016/j.jad.2016.05.040 (2016).

75 Kuroki, N. et al. Fornix integrity and hippocampal volume in male schizophrenic patients. Biological psychiatry 60, 22–31, doi:10.1016/j.biopsych.2005.09.021 (2006).

76 Cheung, V. et al. A diffusion tensor imaging study of structural dysconnectivity in never-medicated, first-episode schizophrenia. Psychological medicine 38, 877–885, doi:10.1017/s0033291707001808 (2008).

77 Bollettini, I. et al. Sterol Regulatory Element Binding Transcription Factor-1 Gene Variation and Medication Load Influence White Matter Structure in Schizophrenia. Neuropsychobiology 71, 112–119, doi:10.1159/000370076 (2015).

78 Doan, N. T. et al. Distinct multivariate brain morphological patterns and their added predictive value with cognitive and polygenic risk scores in mental disorders. NeuroImage. Clinical 15, 719–731, doi:10.1016/j.nicl.2017.06.014 (2017).

79 Nazeri, A. et al. Gray Matter Neuritic Microstructure Deficits in Schizophrenia and Bipolar Disorder. Biological psychiatry, doi:10.1016/j.biopsych.2016.12.005 (2016).

80 Rae, C. L. et al. Deficits in neurite density underlie white matter structure abnormalities in first-episode psychosis. Biological psychiatry, doi:10.1016/j.biopsych.2017.02.008.

81 Lyall, A. E. et al. Greater extracellular free-water in first-episode psychosis predicts better neurocognitive functioning. Molecular psychiatry, doi:10.1038/mp.2017.43 (2017).

82 White, N. S., Leergaard, T. B., D’Arceuil, H., Bjaalie, J. G. & Dale, A. M. Probing tissue microstructure with restriction spectrum imaging: Histological and theoretical validation. Human brain mapping 34, 327–346, doi:10.1002/hbm.21454 (2013).

83 First, M., Spitzer, R. L., Gibbon, M. & Williams, J. B. W. Structured Clinical Interview for DSM-IV Axis I Disoders, Patient Edition (SCID-P). (New York State Psychiatric Institute, 1995).

84 Spitzer, R. L. et al. Utility of a new procedure for diagnosing mental disorders in primary care. The PRIME-MD 1000 study. Jama 272, 1749–1756 (1994).

85 Smith, S. M. et al. Advances in functional and structural MR image analysis and implementation as FSL. NeuroImage 23 Suppl 1, S208–219, doi:10.1016/j.neuroimage.2004.07.051 (2004).

86 Woolrich, M. W. et al. Bayesian analysis of neuroimaging data in FSL. NeuroImage 45, S173–186, doi:10.1016/j.neuroimage.2008.10.055 (2009).

87 Jenkinson, M., Beckmann, C. F., Behrens, T. E., Woolrich, M. W. & Smith, S. M. FSL. NeuroImage 62, 782–790, doi:10.1016/j.neuroimage.2011.09.015 (2012).

88 Andersson, J. L., Skare, S. & Ashburner, J. How to correct susceptibility distortions in spin-echo echo-planar images: application to diffusion tensor imaging. NeuroImage 20, 870–888, doi:10.1016/s1053-8119(03)00336-7 (2003).

89 Andersson, J. L. & Sotiropoulos, S. N. An integrated approach to correction for off-resonance effects and subject movement in diffusion MR imaging. NeuroImage 125, 1063–1078, doi:10.1016/j.neuroimage.2015.10.019 (2016).

90 Andersson, J. L., Graham, M. S., Zsoldos, E. & Sotiropoulos, S. N. Incorporating outlier detection and replacement into a non-parametric framework for movement and distortion correction of diffusion MR images. NeuroImage 141, 556–572, doi:10.1016/j.neuroimage.2016.06.058 (2016).

91 Basser, P. J. & Pierpaoli, C. Microstructural and physiological features of tissues elucidated by quantitative-diffusion-tensor MRI. 1996. Journal of magnetic resonance (San Diego, Calif.: 1997) 213, 560–570, doi:10.1016/j.jmr.2011.09.022 (2011).

92 Roalf, D. R. et al. The impact of quality assurance assessment on diffusion tensor imaging outcomes in a large-scale population-based cohort. NeuroImage 125, 903–919, doi:10.1016/j.neuroimage.2015.10.068 (2016).

93 Liu, Z. et al. Quality Control of Diffusion Weighted Images. Proceedings of SPIE--the International Society for Optical Engineering 7628, doi:10.1117/12.844748 (2010).

94 Roalfa, D. R. et al. The impact of quality assurance assessment on diffusion tensor imaging outcomes in a large-scale population-based cohort. NeuroImage 125, 903–919 (2016).

95 Jenkinson, M., Bannister, P., Brady, M. & Smith, S. Improved optimization for the robust and accurate linear registration and motion correction of brain images. NeuroImage 17, 825–841 (2002).

96 Winkler, A. M., Ridgway, G. R., Webster, M. A., Smith, S. M. & Nichols, T. E. Permutation inference for the general linear model. NeuroImage 92, 381–397, doi:10.1016/j.neuroimage.2014.01.060 (2014).

97 Smith, S. M. & Nichols, T. E. Threshold-free cluster enhancement: addressing problems of smoothing, threshold dependence and localisation in cluster inference. NeuroImage 44, 83–98, doi:10.1016/j.neuroimage.2008.03.061 (2009).

98 R: A language and environment for statistical computing (R Foundation for Statistical Computing, Vienna, Austria, 2014).

99 Davison, A. C. & Hinkley, D. V. Bootstrap Methods and Their Applications. (Cambridge University Press, 1997).

100 Angelo Canty & Ripley, B. D. boot: Bootstrap R (S-Plus) Functions. (2017).

101 Benjamini, Y. & Hochberg, Y. Controlling the False Discovery Rate: A Practical and Powerful Approach to Multiple Testing.. Journal of the Royal Statistical Society. Series B (Methodological), 57, 289–300 (1995).

102 Zeileis, A. & Grothendieck, G. zoo: S3 Infrastructure for Regular and Irregular Time Series. J Journal of Statistical Software 14 (2005).

103 Wickham, H. ggplot2: Elegant Graphics for Data Analysis. (Springer-Verlag New York, 2009).

104 Wakana, S. et al. Reproducibility of quantitative tractography methods applied to cerebral white matter. NeuroImage 36, 630–644, doi:10.1016/j.neuroimage.2007.02.049 (2007).

105 Hua, K. et al. Tract probability maps in stereotaxic spaces: analyses of white matter anatomy and tract-specific quantification. NeuroImage 39, 336–347, doi:10.1016/j.neuroimage.2007.07.053 (2008).

